# Single cell transcriptomics of the human parasite *Schistosoma mansoni* first intra-molluscan stage reveals tentative tegumental and stem cell regulators

**DOI:** 10.1101/2023.05.02.538881

**Authors:** Carmen L. Diaz Soria, Teresa Attenborough, Zhigang Lu, Jennie Graham, Christopher Hall, Sam Thompson, Toby G. R. Andrews, Kate A. Rawlinson, Matthew Berriman, Gabriel Rinaldi

## Abstract

**Background:** Schistosomiasis is a major Neglected Tropical Disease, caused by the infection with blood flukes in the genus *Schistosoma*. To complete the life cycle, the parasite undergoes asexual and sexual reproduction within an intermediate snail host and a definitive mammalian host, respectively. The intra-molluscan phase provides a critical amplification step that ensures a successful transmission. However, the cellular and molecular mechanisms underlying the development of the intra-molluscan stages remain poorly understood.

**Methods:** *S. mansoni* mother sporocysts were dissociated into single cell suspensions, and live cells enriched and sequenced using the single cell 10X Genomics Chromium platform. We defined somatic and stem/germinal cell clusters, identified cell type-enriched Gene Ontology (GO) terms, and predicted transcription factor binding sites for key marker genes.

**Results:** Six cell clusters comprising stem/germinal, two tegument, muscle, neuron, and parenchyma were identified and validated by Fluorescence *in situ* Hybridisation (FISH). GO term analysis predicted key biological processes for each of the clusters. Using the Self-Assembling Manifold (SAM) algorithm, three sub-clusters were identified within the stem/germinal cell population. Furthermore, transcription factor binding sites and putative transcription factors were predicted for stem/germinal and tegument clusters.

**Conclusions:** We report a spatially validated single cell transcriptomic analysis of the first intra-molluscan stage of *S. mansoni.* Key cell regulators were identified, paving the way for future analyses to unveil their role during the parasite development and interaction with its intermediate host.

## Introduction

Schistosomiasis, the infection with parasitic flatworms from the genus *Schistosoma*, remains a major neglected tropical disease with more than 240 million people affected worldwide and more than 700 million at risk of infection in endemic areas^1,2^. To date, a single drug (Praziquantel) is in use; however, this drug is effective only against adult worms, it does not prevent reinfection and drug resistance may be emerging in the field^3^. One approach that may lead to novel control strategies is to gain a better understanding of the mechanisms underlying life cycle progression, including the cells and their transcriptomic signatures across developmental stages^4,5^.

*Schistosoma mansoni* eggs laid by adult worm pairs dwelling in the portal system of the mammalian host traverse the intestinal wall and pass with the faeces to the environment. In contact with freshwater, the eggs hatch free-swimming larvae (miracidia) that seek, infect a suitable snail, and transform into mother sporocysts, the first intra-molluscan developmental stage. Within the mother sporocyst, groups of stem cells (historically termed ‘germinal cells’) start to proliferate and differentiate to develop into a daughter sporocyst^6^. By 5 days post infection (5 dpi), developing daughter sporocysts, initially spherical in shape, start to grow within the brood chamber, elongate and become surrounded by a primitive epithelium derived from the mother sporocyst tegument^6^. By ∼15 dpi they have acquired the definitive vermiform shape containing germinal cells closely packed in the medial part of the body^6^. The mature daughter sporocysts escape from the mother sporocyst, migrate through the snail tissue to the digestive gland area and start to produce whole cercariae from single germinal cells following a second round of embryogenesis^7^. Therefore, from single miracidium hundreds to thousands clonal human-infective cercariae are produced. Extensive knowledge gathered for several decades through detailed histological and electron microscopy-based studies paved the way towards an understanding of the parasite progression within the snail^8–11^. This knowledge can now be scrutinised using current molecular and ‘omics’ technologies^12^. Shining new light on the cellular and molecular basis of this parasite expansion strategy is critical not only to discover novel aspects of trematode developmental biology, but also to reveal targets for control^13^.

Single-cell transcriptome sequencing (scRNA-seq) has been employed to define cellular subtypes by revealing their specific transcriptional signatures. Compared with so-called ‘bulk RNA-seq’ studies of whole organisms or tissues, scRNA-seq has exceptional resolving power, being able to detect genes expressed in just a few cells or with low expression levels, but also reveals the stochastic nature of gene expression in individual cells^14^. Single cell transcriptomics have been used in several systems to understand diverse biological processes, such as cell differentiation, tissue specification and development as well as to define “atlases” cell types^15^ within a given lifecycle stage based on their scRNA-seq profiles. Studies have employed scRNA-seq in *S. mansoni*^5,13^ to generate cell atlases for male and female adult worms^16^, mixed-sex schistosomula^17^, the first intra-mammalian developmental stage, and more recently mixed-sex miracidia^18^. In addition, scRNA-seq has been employed to define and functionally characterise stem cell populations driving the development of both intra-molluscan^19^ and intra-mammalian stages^4^.

Important contributions to our current understanding of the developmental biology of schistosome intra-molluscan stages have been made using ‘bulk RNA-seq’^20^. In particular, two germinal cell lineages with distinct proliferation properties had previously been identified and functionally-characterised by ‘bulk transcriptomics’ and RNAi, respectively^20^. However, there is a critical lack of transcriptomic data and knowledge of gene regulatory networks at the single cell level. Wang and collaborators pioneered the sequencing of 35 sporocyst individual cells, focusing primarily on proliferating stem/germinal cells^19^ but most other cell types of this lifecycle stage remain uncharacterised. In the present study, we have followed an untargeted approach to characterise the individual transcriptomic signatures of more than 600 cells isolated from cultured mother sporocysts. Each of the clusters was spatially validated by fluorescence *in situ* hybridisation (FISH).

Furthermore, we have explored aspects of gene expression regulation, including the prediction of promoter motifs as tentative binding sites for transcription factors, in the stem/germinal and tegumental cell populations. This study contributes to the expansion of the currently scarce number single cell datasets for schistosomes^5^ and reveals key candidate genes involved in the intra-snail developmental phase of this neglected tropical disease pathogen.

## Materials and methods

### Ethics statement

The complete life cycle of *Schistosoma mansoni* (NMRI strain) was maintained at the Wellcome Sanger Institute (WSI). All the animal regulated procedures were conducted under Home Office Project Licence No. P77E8A062 held by GR. All the protocols were presented and approved by the Animal Welfare and Ethical Review Body (AWERB) of the WSI. The AWERB is constituted as required by the UK Animals (Scientific Procedures) Act 1986 Amendment Regulations 2012.

### Parasite material

On a monthly basis, TO outbred mice were infected as described^21^. The animals were culled six weeks after infection and livers were removed and processed for egg isolation^22,23^. Parasite material was prepared as previously described^23^. The livers were finely minced and digested overnight at 37°C with gentle agitation in 1x PBS containing 0.5% of *Clostridium histolyticum* collagenase (Sigma), 200 U/ml penicillin, 200 μg/ml streptomycin and 500 ng/ml amphotericin B (ThermoFisher Scientific). The resulting mixture was filtered through sterile 250 μm and 150 μm sieves and subjected to a Percoll-sucrose gradient to purify the eggs, which were further washed three times in 1x PBS and 200 U/ml penicillin, 200 μg/ml streptomycin and 500 ng/ml amphotericin B. The eggs were transferred to sterile water to induce hatching under light for at least 3 hours, collecting miracidia and replacing the water every ∼30 min. The miracidia were incubated for 30 min on ice, centrifuged at 800 g, 4°C for 15 min, resuspended in sporocyst medium (MEMSE-J, 10% Fetal Bovine Serum, 10mM Hepes, 100 U/ml penicillin, 100 μg/ml streptomycin – all reagents obtained from ThermoFisher Scientific) and cultured in a hypoxia chamber, with a gas mixture of 1% O_2_, 3% CO_2_ and 96% N_2_, at 28 °C for 5 days^22,23^.

### Sporocyst cell number estimation by nuclear segmentation

*In-vitro* transformed mother sporocysts were cultured for 5 days (D5 sporocysts) and collected, washed three times by centrifugation, 400 g for 5 min, 1x PBS and fixed overnight in 4% methanol-free paraformaldehyde (Pierce^TM^) in 1x PBS at 4°C. The parasites were washed three times in 1x PBS as above, resuspended in mounting media containing 4’,6’-diamidino-2-phenylindole (DAPI) (Fluoromount-G™ Mounting Medium, with DAPI, Invitrogen), incubated overnight at 4°C, and mounted on microscope slides. Z-stacks and maximum intensity projections were taken with a Leica SP8 confocal microscope. Nuclear segmentation was performed using the machine learning platform Ilastik (version 1.3.3)^24^. To enable the detection of nuclei with heterogeneous DAPI staining, pre-processing steps were first performed in ImageJ^25^, which involved applying image down-scaling, a Gaussian filter (2 pixel radius), and a median filter (2 pixel radius). A binary pixel classification was then trained in Ilastik^24^ using the auto-context pipeline, which includes two rounds of pixel classification. Training was performed on two small regions of interest from the dataset, representative of the total diversity of nuclear staining quality and intensity. Hysteresis thresholding was then used to segment the binary pixel classification map into distinct objects representing individual nuclei, with a core threshold of 0.93 and final threshold of 0.5. To further improve nuclear segmentation, thresholded images were imported to ImageJ using the Ilastik plugin, and a 3D distance-transform watershed was applied from the MorphoLibJ package^26^.

### Single-cell tissue dissociation and Fluorescence-activated Cell Sorting (FACS)

D5 sporocysts were collected and processed for tissue dissociation as described^17^. Briefly, ∼5,000 D5 sporocysts were collected in 15-ml tubes and digested for 30 min in an Innova 4430 incubator with agitation at 300 rpm at 37°C, using a digestion solution of 750 μg/ml Liberase DL (Roche) in 1x PBS supplemented with 20% heat inactivated FBS. The resulting cell suspension was successively passed through 70μm and 40μm cells strainers, centrifuged at 300 g for 5 mins and resuspended in cold 1x PBS supplemented with 20% FBS. The cells were co-stained with 0.5 μg/ml of Fluorescein Diacetate (FDA; Sigma) and 1 μg/ml of Propidium Iodide (PI; Sigma) to label live and dead/dying cells, respectively. Live cells were enriched using BD Influx™ cell sorter (Becton Dickinson, NJ). It took 2-3 hours from the enzymatic digestion to the generation of single-cell suspensions ready for loading on the 10X Genomics Chromium platform.

## 10X Genomics library preparation and sequencing

Suspensions of ∼ 500 cells/μl live single-cells from mother sporocysts were loaded according to the standard protocol of the Chromium single-cell 3’ kit to capture ∼ 7,000 cells per reaction (V2 chemistry). Thereafter, single-cell libraries were prepared and sequenced on an Illumina Hiseq4000 (custom read length: 26 bp read 1, 98 bp read 2, 8 bp index 1, 0 bp index 2 using the 75 bp PE kit), one sequencing lane per sample. Four biological replicates, i.e., 4 independent sporocyst batches generated from eggs collected in different occasions, were sequenced and analysed. All raw sequence data is deposited in the ENA under the project accession ERP137194 and sample accession numbers: ERS11891013, ERS11891016, ERS11891015, ERS11891014 (**Supplementary Table S1**).

### Mapping and single-cell RNA-seq quantification

The reference FASTA sequence for *S. mansoni* genome (version 9) was downloaded from WormBase Parasite release 17^27,28^ and gene annotations obtained from an intermediate version 9 stage before the WBPS release 17 (gtf accessible upon request from the authors). We followed previously described protocols with minor modifications^17^. In brief, scRNA-seq data were mapped to the *S. mansoni* reference genome using Cell Ranger (v 6.0.1), with default parameters.

Approximately 62.2% of sequenced reads (average over four samples) mapped confidently to the transcriptome with a median UMI count per cell ranging from 973 to 1,198 and a total of 857 cells detected (**Supplementary Table S1**).

### Quality control, cell clustering and marker identification

The Seurat package (version 4.1.1)^29^ (https://satijalab.org/seurat/) was used to analyse the filtered counts matrix produced by Cell Ranger. A maximum mitochondrial read threshold was set at 3% per cell. Analysis and quality control (QC) was performed according to the standard Seurat workflow using LogNorm (https://satijalab.org/seurat/)^16^. Following QC, 601 cells remained for downstream analyses. A combination of Seurat’s JackStraw and Elbowplot were used to select the first 12 principal components for clustering cells. Clusters were generated using FindNeighbors and FindClusters, and clustree was used to choose the resolution for stable clustering. We manually segmented one cluster into two based on clear expression of neural markers, resulting in the identification of six cell clusters in total. Top markers for each cell cluster were identified using Seurat’s FindAllMarkers function (test.use □ = □ “roc”, only.pos = TRUE, return.thresh = 0), projecting cluster annotation from schistosomula scRNA-seq data^17^, and comparing to top markers curated from the literature^13,19^.

The data were further analysed using the self-assembling manifold (SAM) algorithm^30^ (available at https://github.com/atarashansky/self-assembling-manifold) to assess cell cluster structure. The raw count data from the cells passing QC were exported from the Seurat object and imported to a SAM object, pre-processed and the SAM algorithm run using the default parameters (using Python version 3.8.5). To analyse the stem cells specifically, the above was repeated but only exporting cells from the annotated stem cell cluster (119 cells). After running the SAM algorithm to find topology, leiden clustering was overlaid (param=0.8). Scanpy^31^ (version 1.8.2) was then used to extract summary QC information and the variable genes that characterised the three SAM clusters (rank_genes_groups). The cells were grouped by leiden cluster to rank the genes that characterise each group (method=wilcoxon, corr.method=benjamini-hochberg).

### Gene Ontology (GO) and motif analysis

To create specific marker gene lists for Gene Ontology (GO) term analysis, the Seurat marker gene list was filtered for each cluster such that the remaining markers had an AUC score ≥ 0.7 (**Supplementary Table S4)**. For the motif analyses (below), this gene list was further filtered for specificity so each gene had to be detected in ≥ 70% cells in the considered cluster, and ≤ 30% of cells in the other clusters. GO analysis was also performed with a more permissive marker gene list filtering of AUC score ≥ 0.6, and results are available in **Supplementary Table S5**.

### GO enrichment analysis

GO term enrichment was performed using the weight01 method provided in topGO v2.46.0 (available at http://bioconductor.org/packages/release/bioc/html/topGO.html) for the Biological Process (BP) and Molecular Function (MF) aspects of the ontology. Analysis was restricted to terms with a node size of ≥ 5. Fisher’s exact test was applied to assess the significance of overrepresented terms compared with all expressed genes. The threshold was set as FDR < 0.05. Analysis of GO term enrichment for BP was also performed for comparison with other *S. mansoni* developmental stages for which single-cell transcriptomic data are available. GO term analysis tables from previously published developmental stages were extracted from supplementary tables^16,18^ or manually^17^. RVenn (v1.1.0) was used to find and visualise overlapping GO terms (by GO ID) in sporocyst tegument clusters with tegument GO terms from miracidia^18^, schistosomula^17^, and adults^16^.

### Motif analysis in the promoter region of marker genes

The sequences of promoter regions (defined as 1kb upstream of annotated transcriptional start sites or the intervening distance to the upstream gene if < 1kb) of marker genes were extracted based on the *S. mansoni* v9 annotation and fed to the MEME Suite tool XSTREME (v5.4.1; https://meme-suite.org/meme/tools/xstreme)^32^. We used shuffled input sequences with default Markov order as the control. In addition, the non-redundant transcription factor (TF) binding profiles (Position Frequency Matrix) for nematodes were obtained from the JASPAR 2022 database^33^ and supplied to XSTREME. A default threshold *E*-value < 0.05 was applied to select enriched motifs in the promoter regions and *E*-value < 1 for matching the motif to JASPAR TF binding sites using the Tomtom tool. Based on the discovered transcription factor binding sites (TFBS), peptide sequences of corresponding transcription factors in *C. elegans* were retrieved from WormBase^34^ and aligned to *S. mansoni* proteins using BlastP with *E*-value < 1e-5; *C. elegans* was chosen as having a particularly well-curated set of TFBS and TFs. The top 10 *S. mansoni* protein hits for each *C. elegans* TF (including isoforms) were checked for their TF identity based on KEGG^35^ ortholog search using the GHOSTX program and BBH method, and their binding profiles were validated again using the Jaspar Profile Inference tool (https://jaspar.genereg.net/inference).

### Protein-protein interaction network analysis

To predict protein-protein interactions within the stem/germinal subcluster top markers, we used the online tool STRING^36^ (https://string-db.org). In STRINGdb, interaction networks were constructed using evidence sources as edges and a minimum interaction score of 0.9 for the top 50 marker genes from the SAM analysis.

### Fluorescence *in situ* hybridisation

To spatially validate the main cell populations, we used fluorescence *in situ* hybridisation (FISH) following the third-generation *in situ* hybridisation chain reaction (HCR) approach^37^. Buffers, hairpins and probes against top marker gene transcripts for tegument (*micro-exon gene 6* or *MEG-6;* Smp_163710), muscle (*myosin heavy chain;* Smp_085540), stem/germinal cell (*histone H2A;* Smp_086860), parenchyma (*hypothetical protein;* Smp_318890) and neuronal (*neuroendocrine protein 7b2; Smp_*073270) cell clusters were purchased from Molecular Instruments (Los Angeles, California, USA) (**Supplementary Table S2**). D5 sporocysts were processed for HCR as described^17,18,38^, with minor modifications. In brief, the D5 sporocysts were collected, washed once in sporocyst media (above) by centrifugation at 500 g, 5 min at room temperature (RT), and fixed in 4% PFA in PBSTw (1x PBS + 0.1% Tween) at RT for 30 min with gentle rocking. The fixed sporocysts were transferred into an incubation basket (Intavis, 35µm mesh) in a 24-well plate, rinsed 5 times for 5 min in ∼750 μl of PBSTw, and incubated in 750 μl of 1x PBSTw containing 2 μl of Proteinase K (20 mg/ml) (ThermoFisher Scientific) at RT with no agitation. Thereafter, the sporocysts were rinsed twice in ∼750 μl of PBSTw for 5 min at RT, incubated in 500 μl of PBSTw with 0.2 mg/ml glycine on ice for 15 min and rinsed twice as above. Thereafter, the parasites were distributed into different baskets depending on the number of experimental groups to test (up to two probes were multiplexed per experimental group), and incubated in 750 μl of 1:1 PBSTw: hybridisation buffer for 5 min at RT. The 1:1 PBSTw: hybridisation buffer solution was replaced with 750 μl of hybridisation buffer, and parasites placed in a humidified chamber and incubated at 37 °C for 1 h (i.e., pre-hybridisation step). During the incubation, 1 to 2 μl of indicated HCR probe was added to 650 μl of hybridisation buffer and equilibrated to 37°C. After the pre-hybridisation step, the hybridisation buffer was replaced with the hybridisation buffer containing the HCR probe(s), and parasites incubated at 37°C in the humidified chamber overnight. The parasites were rinsed four times for 15 min in probe wash buffer, previously equilibrated at 37°C, followed by four washes in filtered 5x SSCTw (5 x SSC buffer + 0.1% Tween) at RT on gentle rocker and incubated in 600 μl of amplification buffer, previously equilibrated at RT, for 30 min at RT on gentle rocker. During this pre-amplification step, 10 μl of corresponding hairpin for each probe were incubated at 95 °C for 90 sec, cooled to RT, in the dark for 30 min and added to 600 μl of amplification buffer. Thereafter, the amplification buffer was replaced with amplification buffer containing the hairpin mix, and the plate was incubated in the dark, at RT overnight. The larvae were rinsed in 5 x SSCTw, three times for 5 min followed by a long wash of 30 to 60 min on gentle rocker in the dark. In some experiments, the sporocysts were incubated in phalloidin (1 μl of phalloidin diluted in 2 ml of 5x SSCTw) and incubated in the dark for ∼60 min followed by three washes of 5 min each and four washes of 30 min each in 5 x SSCTw. After the washes, the parasites were incubated in mounting media containing DAPI (Fluoromount-G™ Mounting Medium, with DAPI, Invitrogen), incubated overnight at 4°C, and mounted on microscope slides. For each HCR experiment, controls of hairpin with no probe (i.e., control for signal specificity), and no probe no hairpin (i.e., control for autofluorescence) were included. Z-stacks and maximum intensity projections were taken with a Leica SP8 confocal microscope. FISH experiments were performed at least twice (≥ 2 different batches of sporocysts collected from different groups of infected mice) for each HCR probe. The number of screened parasites (*n*) are indicated in the Figure legends.

## Results

### Six cell populations identified in the *Schistosoma mansoni* mother sporocyst

Freshly collected *S. mansoni* miracidia were transferred into sporocyst media to induce transformation into mother sporocysts. Within the first ∼16 h most parasites have shed the cilia plates and their tegument has been remodelled^8^; however, parasite *in-vitro* development is not synchronous. Therefore, we decided to culture the mother sporocysts for 5 days (named ‘D5 sporocysts’) to facilitate the complete transformation of >95% of the parasites^10,39^ (**Supplementary Figure S1**; **Figure 1A**). The D5 sporocysts were collected and processed following the dissociation protocol previously used for schistosomula^17^, and live cells enriched and quantified using FACS. The droplet-based 10X Genomics Chromium platform was used to generate transcriptome-sequencing data from a total of 601 cells after applying quality-control filters. With the assistance of a high-resolution nuclei quantification protocol, based on a machine-learning imaging platform^24^, we estimated that a D5 sporocyst comprises an average of 169 nuclei (range: 112-254) (**Supplementary Figure S2**). Therefore, the number of quality-controlled cells theoretically represents >3.5x coverage of all cells in a single D5 mother sporocyst.

**Figure 1.**
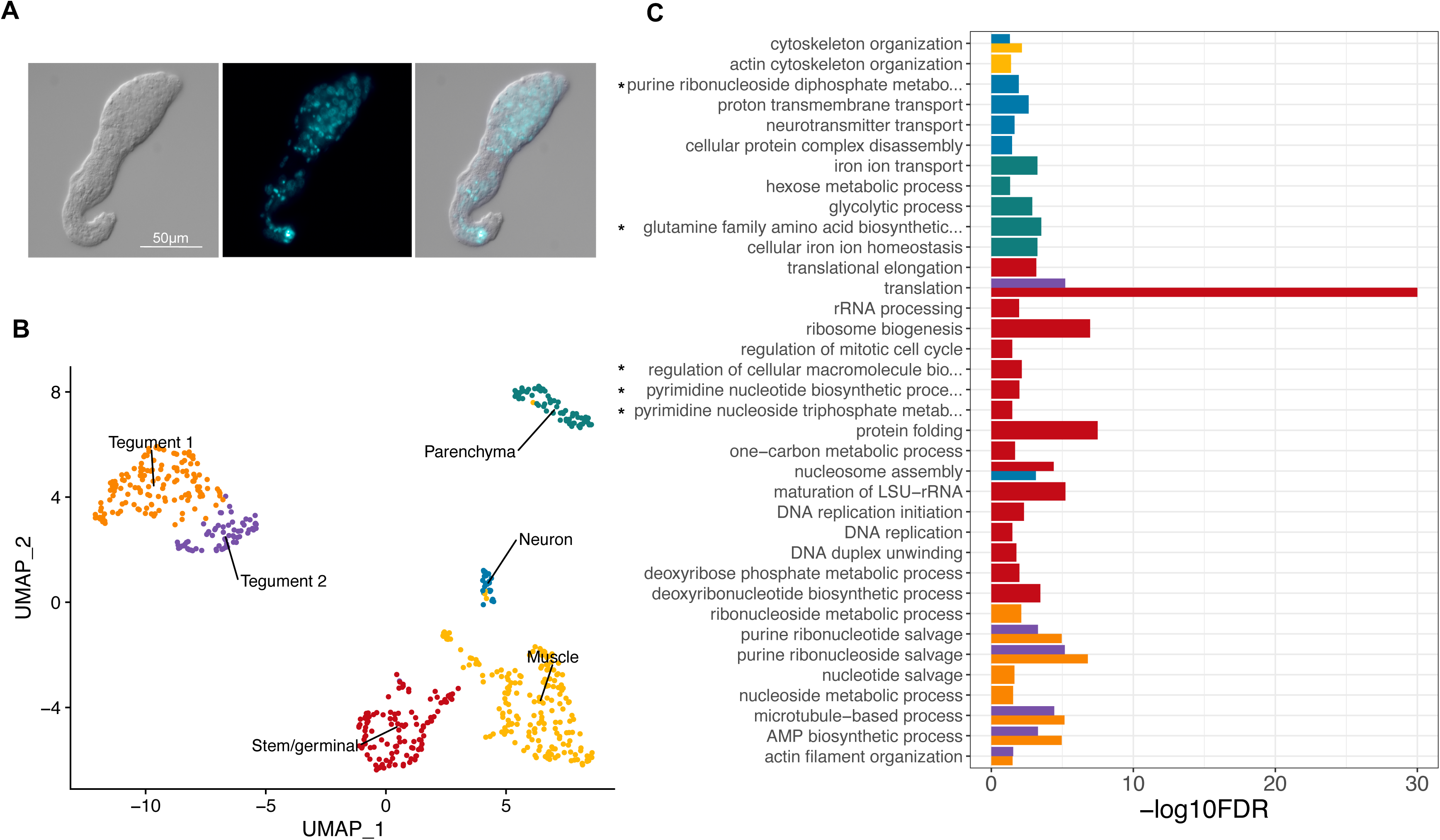
Six distinct cell populations identified in D5 mother sporocysts. **A.** Representative picture of D5 sporocyst in DIC bright field (left), DAPI-fluorescent field (centre) and merged (right). Scale bar: 50 μm. **B.** Uniform Manifold Approximation and Projection (UMAP) representation of 601 single cells from D5 sporocysts. The cell clusters are coloured and labelled as indicated. The list of the Seurat marker genes in all cell clusters is provided in **Supplementary Table S3**. **C.** Gene ontology (GO) enrichment analysis for biological processes only (marker genes with minimum AUC=0.7, and GO terms supported by 172 genes), for top marker genes in each indicated colour-coded cell cluster (as shown in B). Only statistically significant GO enriched biological processes are depicted (-log10 (FDR<0.05)). FDR: False Discovery Rate. Full names of GO terms indicated with *: purine ribonucleoside diphosphate metabolic process (GO:0009179), glutamine family amino acid biosynthetic process (GO:0009084), regulation of cellular macromolecule biosynthetic process (GO:2000112), pyrimidine nucleotide biosynthetic process (GO:0006221), pyrimidine nucleoside triphosphate metabolic process (GO:0009147). Full data provided in **Supplementary Table S4** and **Supplementary Table S5**, for non-specificity filtering analysis with AUC=0.7 and AUC=0.6, respectively.

Based on top markers identified using Seurat, annotation from schistosomula single cell data and genes curated from the literature, we identified six discrete cell populations (**Supplementary Table S3**); Tegument-1 (138 cells), Tegument-2 (66 cells), Muscle (189 cells), Stem/germinal (119 cells), Parenchyma (66 cells) and Neuron (23 cells) (**Figure 1B**). To further explore the biological processes in which the cells of each cluster are involved, we examined over-represented Gene Ontology terms using TopGO (**Figure 1C**). Within each cluster, there were clear examples of single genes with annotated roles that are statistically enriched due to their rarity in the genome (**Supplementary Tables S4 and S5**). Although many of these conformed to expectations — for instance, ‘neuropeptide signalling’ is enriched in the Neuron cluster due to the expression of the known marker gene *7B2* — we focussed on over-represented annotation supported by multiple genes. In the case of the Neuron cluster, this highlighted ‘neurotransmitter transport’, as well as less expected terms ‘nucleosome assembly’ and ‘proton transmembrane transport. In Muscle, ‘cytoskeletal organisation’ was clearly enriched and as expected, Parenchyma showed enrichment for several aspects of metabolism including, ‘iron homeostasis’, ‘amino acid metabolism’ and ‘glycolysis’. In both Tegument clusters, ‘purine ribonucleotide/side salvage’ was highly enriched, as was ‘microtubule-based process’ due to the expression of 6-8 dyneins and the dynein-domain protein SmTAL2. In Tegument-2, genes involved in translation were also over-represented (**Figure 1C**; **Supplementary Tables S4 and S5**). The unexpected overrepresentation of biological processes associated with ‘nucleotide/side metabolism’ in the sporocyst tegument prompted us to compare among the most significantly represented GO terms in the tegument of developmental stages for which single-cell transcriptomic data are available^16–18^. The comparative analysis among the miracidium^18^, schistosomulum^17^, adult^16^ and sporocyst showed not only an expected GO term present across all stages (i.e., ‘microtubule-based process’), but also specific biological processes enriched only in the sporocyst tegument (**Supplementary Figure S3**). Expectedly, the Stem/germinal cell cluster was highly enriched for genes involved in DNA replication, ribogenesis and translation. These results suggest that cluster-specific gene products display distinct functions in different molecular pathways, and that the identified cell populations may be involved in different biological processes (**Figure 1C**; **Supplementary Tables S4 and S5**).

To spatially validate the predicted cell clusters, we defined highly specific cluster-defining marker genes (**Figure 2A**; **Supplementary Table S6**), for which Fluorescence *in situ* Hybridization (FISH) probes were generated (**Supplementary Table S2**). We identified cells expressing the Muscle-specific marker myosin heavy chain, location of which correlated with actin filaments following the anterior-posterior axis of the sporocyst (**Figure 2B**; **Supplementary Figure S4A, S4B**). Whilst a Neuron-specific marker was expressed across a handful scattered cells in the mid region of the parasite (**Figure 2B**; **Supplementary Figure S4C, S4D**), cells expressing the Stem/germinal cluster-specific marker histone H2A were mainly located in clusters towards one pole (**Figure 2C**). As stem cells are located in the posterior half of the miracidium larva^18^, we hypothesise this is the same in the mother sporocyst, so the Stem/germinal cell markers are thus likely to be highlighting the posterior end. In addition, a few individual Stem/germinal cells were located in the medial region towards the surface of the animal (**Supplementary Figure S4E, S4F**). The two Tegumental cell clusters that highly expressed the micro-exon gene 6 (*MEG-6*), were spatially-validated by a strong FISH signal from cells lining the surface of the parasites (**Figure 2D**; **Supplementary Figure S5**). The Parenchyma cluster cells were identified by the cluster specific expression of Smp_318890 (encoding a hypothetical protein). Even though this marker was expressed in <50% of the cells (**Figure 2A**), it showed a high and specific expression in Parenchyma cells. These cells seem to be distributed throughout the whole parasite body with a tendency towards the posterior pole (Figure 2E; **Supplementary Figure S4G, S4H**). The parenchymal cells showed clear anterior-posterior cytoplasmic projections containing Smp_318890 transcripts (**Figure 2E**).

**Figure 2.**
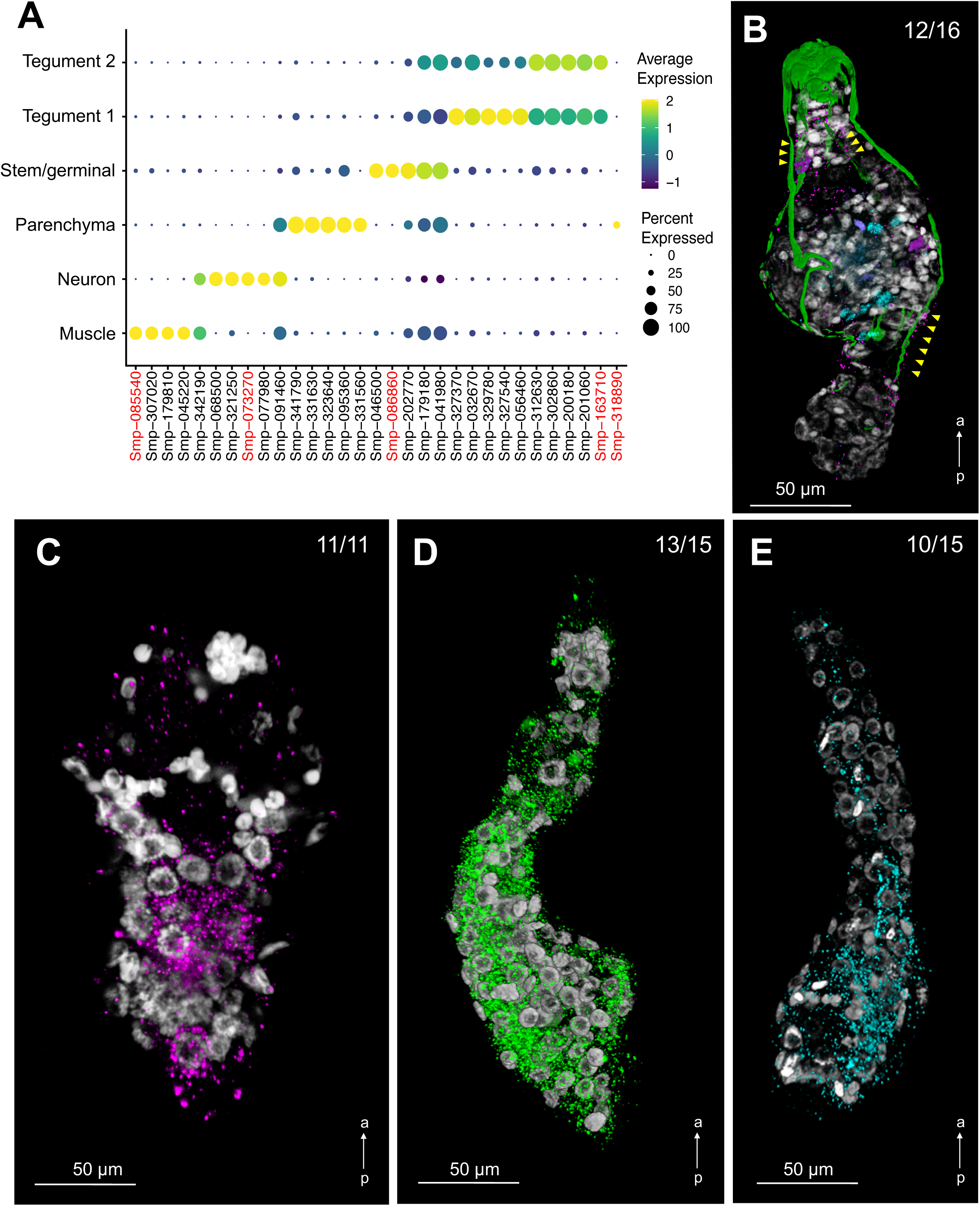
Cell clusters spatially validated by fluorescence *in situ* hybridization (FISH). **A.** Dot plot showing the expression of the top 5 markers identified for each cell cluster. The average gene expression level for each marker is represented by a colour gradient from dark blue (low expression) to bright yellow (high expression). The circle sizes indicate the percentage of cells in each indicated cluster. FISH probes for the following cluster-specific markers (highlighted in red) were used for spatial validation: Pan-tegument, *micro-exon gene 6* or *MEG-6 (*Smp_163710); Muscle, *myosin heavy chain (*Smp_085540); Stem/Germinal, *histone H2A (*Smp_086860); Parenchyma, hypothetical protein (Smp_318890); and Neuron, *neuroendocrine protei*n *7b2* (Smp_073270). Full data for the indicated top markers are provided in **Supplementary Table S6**. **B.** Double FISH with *myosin heavy chain* (Smp_085540; magenta) and *neuroendocrine protein 7b2* (Smp_073270; cyan) probes identified muscle and neuron cell clusters, respectively. Phalloidin-stained actin filaments are shown in green and DAPI staining in grey. Yellow arrowheads indicate co-localisation of *myosin heavy chain* and actin filaments. *n* = 16 parasites. **C.** Localisation of the stem/germinal cells using FISH with *histone H2A* (Smp_086860; magenta). DAPI staining in grey. *n* = 11 parasites. **D.** Localisation of tegumental cells using FISH with *MEG-6* (Smp_163710; green). DAPI staining in grey. *n* = 15 parasites. **E.** Localisation of parenchyma cells using FISH with *hypothetical protein* (Smp_318890; cyan). DAPI staining in grey. *n* = 15 parasites. Scale bar: 50 μm, a←p: anterior-posterior axis (Panels B-E).

### Stem cell heterogeneity revealed by self-assembling manifold algorithm

Using the self-assembling manifold (SAM) algorithm^30^, the 119 stem/germinal cells were further analysed. Three discrete subclusters (clusters 0, 1 and 2) with distinct transcriptional profiles were identified (**Figures 3A, 3B**; **Supplementary Tables S7, S8**). We used Scanpy^31^ (rank_genes_groups) to rank genes that characterise each of the stem cell subclusters (**Supplementary Table S9**) and the five top-ranked genes for each cluster were used for visualisation. STRING^36^ was used to predict protein-protein interactions among the top 50 ranked genes within each subcluster (**Supplementary Table S10**). The analysis of subcluster 0 showed a network with strong connectivity of inferred interactions among the top marker genes (**Figures 3C**). The top STRING terms for subcluster 0, across all categories, are related to ribosomes and translation (**Supplementary Table S10)**. In contrast, weak connectivity was found between genes expressed in subclusters 1 and 2, with no network, far fewer STRING terms, and weaker statistical support (**Supplementary Table S10**).

**Figure 3.**
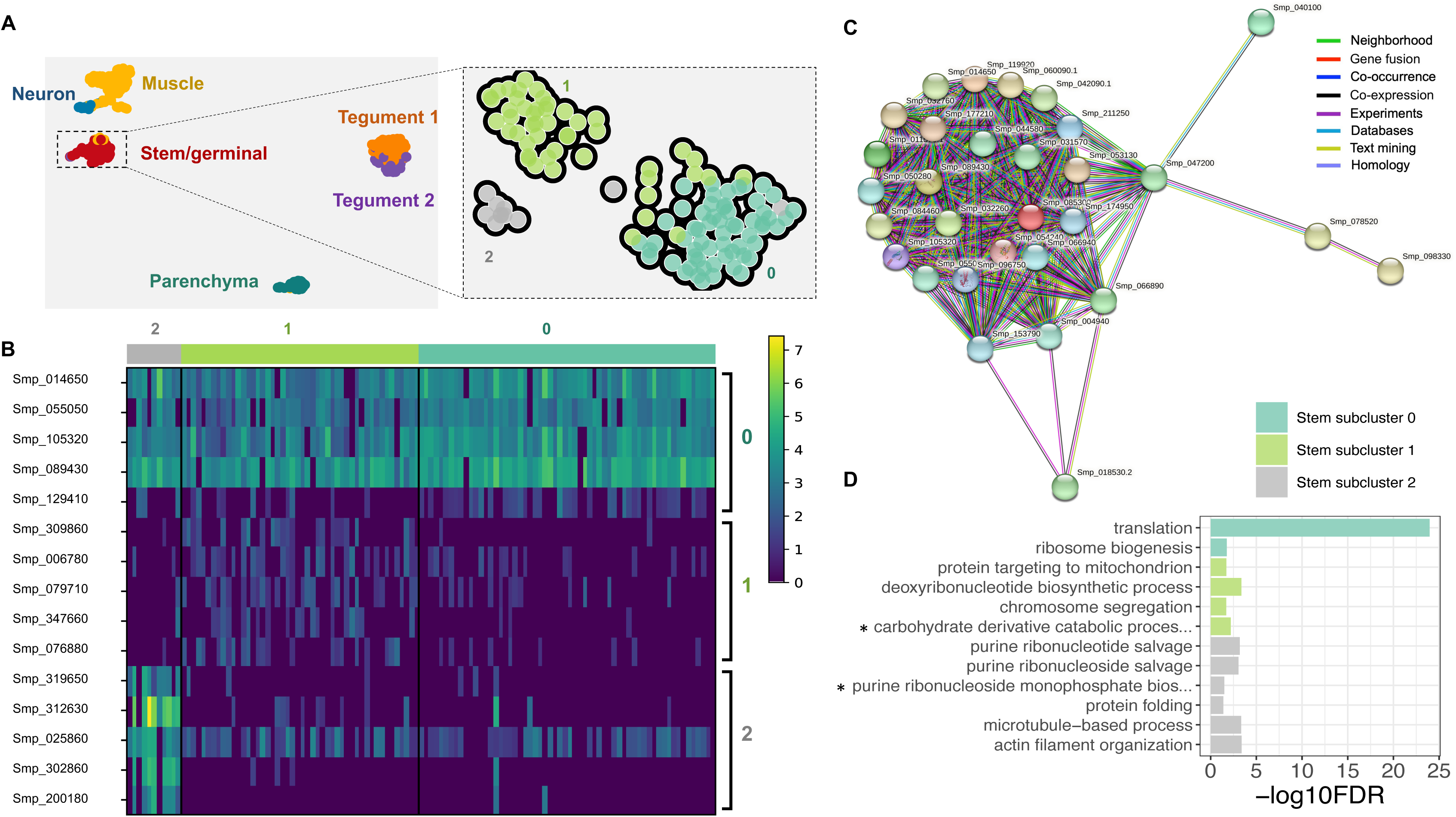
Stem/germinal cell sub-clusters. **A**. Clustering of the sporocyst data using the self-assembling manifold (SAM) algorithm for all cells (left) or the Stem/germinal cluster only (right). The SAM algorithm with Leiden clustering identified three stem/germinal subclusters (0, 1 and 2). The lists of SAM topology genes for all cell clusters or stem/germinal cell cluster only are provided in **Supplementary Table S7** and **Supplementary Table S8**, respectively. **B.** Heatmap of expression of the top 5 marker genes identified in each of the three stem/germinal subclusters. The average gene expression level for each marker is represented by a colour gradient from dark blue (low expression) to bright yellow (high expression). The full list of top marker genes identified by Scanpy in the three SAM stem cell subclusters is provided in **Supplementary Table S9**. **C.** Interaction network analysis by STRINGdb for stem/germinal subcluster 0. The coloured nodes of the network represent proteins (genes ID for each protein are indicated, and all splice isoforms or post-translational modifications for each protein are collapsed). The coloured edges indicate sources of the interaction evidence as described. The full list of all enriched String terms for the top 50 markers of each stem/germinal sub-cluster is provided in **Supplementary Table S10**. **D**. Stem/germinal subcluster gene ontology (GO) enrichment analysis in the category biological process, for top marker genes in each indicated colour-coded cell cluster. Only statistically significant GO enriched biological processes are depicted (-log10 (FDR<0.05)), and only terms supported by 172 genes are shown. FDR: False discovery rate. Full names of GO terms indicated with *: carbohydrate derivative catabolic process (GO:1901136), purine ribonucleoside monophosphate biosynthetic process (GO:0009168). Full data provided in **Supplementary Table S11**.

UsingTopGO we further explored the biological roles of cells within each stem/germinal subcluster by examining over-represented Gene Ontology terms. Related annotations from multiple genes within each cluster were evident; for example, and consistent with the STRING findings, subcluster 0 was highly enriched for biological processes ‘translation’ and ‘ribosome biogenesis’ (**Figures 3D**; **Supplementary Table S11**). Statistically significant biological processes, supported by a minimum of two genes, were identified for the other two subclusters. In subcluster 1, ‘deoxyribonucleotide biosynthesis’ and ‘chromosome segregation’ suggest active DNA synthesis and cell division (**Supplementary Table S11**). In subcluster 2, ‘actin filament organisation’ and ‘microtubule-based process’, supported by three and six genes, respectively, as well as ‘purine ribonucleoside salvage’, ‘purine ribonucleotide salvage’ and ‘purine ribonucleoside monophosphate biosynthetic process’ indicate considerable similarity to the tegumental gene clusters described above (**Supplementary Table S11**).

Previously, three stem cell populations in mother sporocysts^19^ were identified as *kappa*, *delta* and *phi* based on the differential expression of seven marker genes (*klf*, *nanos-2*, *fgfrA fgfrB*, *p53*, *zfp-1*, and *hesl*). Although these markers were expressed, we were not able to use them to unambiguously assign the Stem/germinal cell subclusters identified by SAM to the *kappa*, *delta* and *phi* cell populations (**Supplementary Figure S6**; **Figure S7**; **Figure S8**).

### Promoter motifs and transcription factor binding sites enriched in two major cell populations

To further investigate gene expression regulation in the Stem/germinal cell population, we searched for putative TFBSs. The Tegument-1 cell cluster was included as a control because it belongs to a somatic/differentiated cell population with a similar number of cells as the Stem/germinal cell population. Motif analysis was performed for marker genes specific to the Stem/germinal cluster (12 genes; **Figure 4A**) and the Tegument-1 cluster (49 genes; **Supplementary Figure S9A**). We identified five enriched motifs in the promoter region (i.e., 1kb upstream the TSS) of Stem/germinal cell marker genes, and ten in those of tegumental cells (**Figure 4B**; **Supplementary Figure S9B**; **Supplementary Table S12**). Interestingly, nearly all enriched motifs had ≥1 significant match(s) to known TFBSs in the model worm *C. elegans* based on JASPAR 2022 database, and no overlapping motifs were identified between the two analysed cell clusters (**Supplementary Table S13**). Within both the Stem/germinal (**Figure 4C**) and Tegument-1 cell clusters (**Supplementary Figure S9C**), most marker genes have binding sites for multiple transcription factors (TFs). Notably, from the 12 Stem/germinal cell marker genes, 11 contain the motif S-STREME-1, that is similar to the binding site for *ceh-22*, a homeobox gene and orthologue of the human *NKX2*-2 gene (**Figure 4D**). A further eight genes share the binding motif for *pha-4* (S-STREME-5), a forkhead/winged helix factor, and eight genes share a motif (S-STREME-4) similar to the TFBSs for both *ceh-22* and *vab-7* homeobox genes (**Figure 4C**). Six genes share S-STREME-2, which is similar to the *unc-30* homeobox binding motif, and three genes share S-STREME-3, similar to the binding motif for the transcription factor *sma-4* (SMAD/NF-1 DNA-binding domain factors; **Supplementary Table S13**). Detailed information of motif locations within the promoter region of stem/germinal and tegument 1 marker genes is provided in **Supplementary Table S14**.

**Figure 4.**
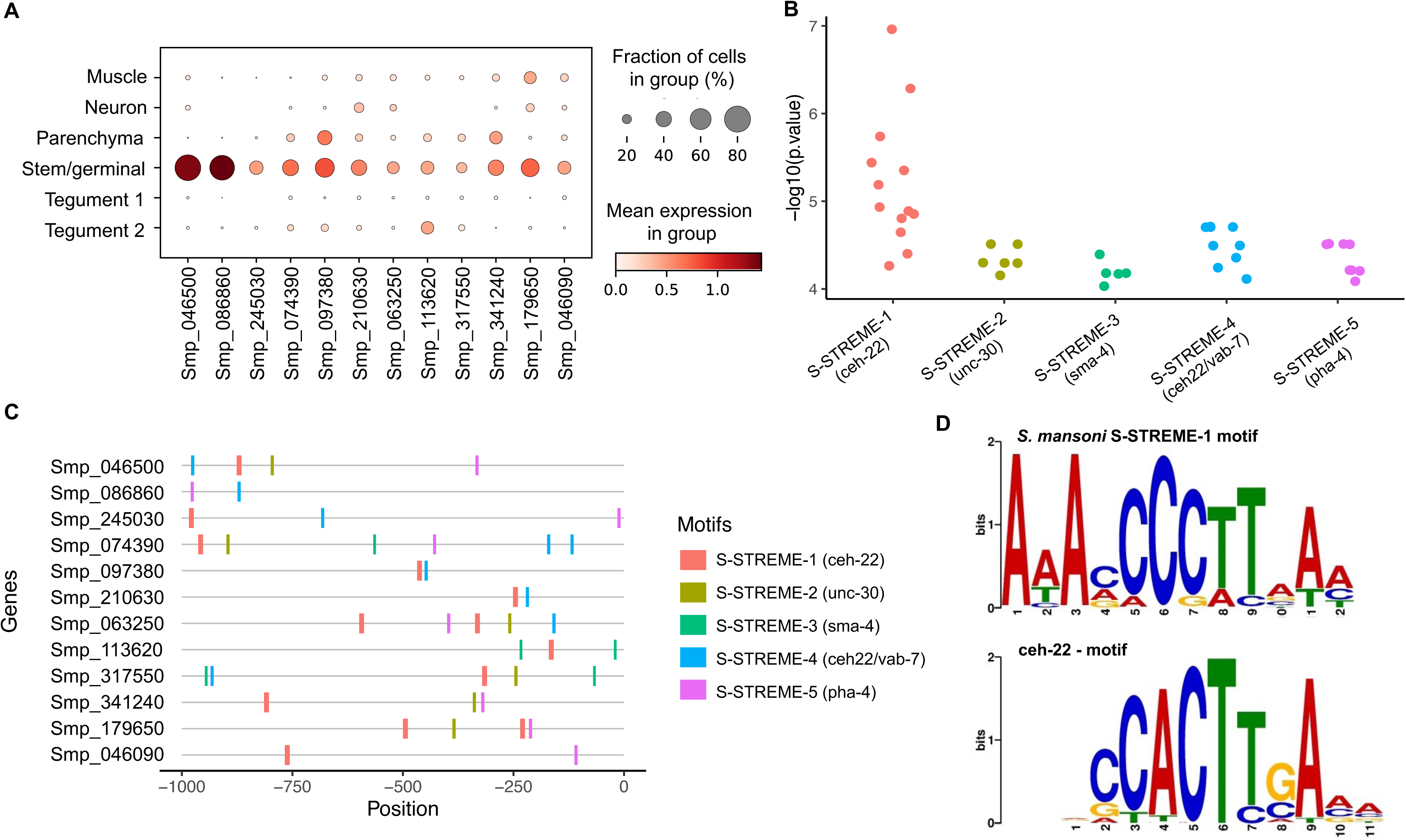
Promoter motif and transcription factor binding sites in stem/germinal cells. **A**. Dot plot showing the expression level of the 12 Stem/germinal cell cluster-specific marker genes used for the analysis. Fraction of cells (%) and mean expression are indicated. The average gene expression level for each marker is represented by a colour gradient from white (low expression) to dark red (high expression). **B**. Distribution of the –log10(p values) for the top 5 ranked motifs identified in the 12 stem/germinal cell marker genes. The x-axis indicates motif names from XSTREME and significant match (p < 0.05) to known Transcription Factors Binding Sites (TFBSs) in the JASPAR 2022 nematode dataset (https://jaspar.genereg.net/downloads/). The y-axis represents log-transformed p values of each motif site shown in C. Full data provided in **Supplementary Table S12** and **Supplementary Table S13. C.** Predicted position distribution of the top 5 ranked motifs along the promoter region of the stem/germinal cell marker genes. The promoter region was taken as 1 kb upstream of the Transcription Start Site (TSS). Full data provided in **Supplementary Table S14**. **D.** TF binding motifs found enriched in the promoter region of stem/germinal cell cluster marker genes. *Schistosoma mansoni* enriched motif named S-STREME-1 with the sequence: 1-AAAMCCCTTAAM (top) found in 11 of the 12 stem/germinal cell cluster marker genes with significant match to the binding site MA0264.1 (MA0264.1.ceh-22) for *C. elegans* ceh-22 (bottom) in the JASPAR database (https://jaspar.genereg.net/). The height of the letter in the motif scheme represents the frequency of the nucleotide observed in each indicated position.

Using the above TFs from *C. elegans*, we identified putative *S. mansoni* orthologs. The putative *S. mansoni* TFs were examined by KEGG ortholog search, and the binding profiles were further confirmed for some candidates using the Jaspar Profile Inference tool^33^. This screening produced a list of 12 candidate *S. mansoni* TF genes (**Supplementary Table S15**): one ortholog of *sma-4* (Smp_033950); two orthologs of *ceh-22* (Smp_027990 and Smp_186930), *pha-4* (FoxA) (Smp_331700 and Smp_158750) and *unc-30* (Smp_124010 and Smp_163140); and five orthologs of *vab-7* (Smp_147640, Smp_138140, Smp_347890, Smp_308310, Smp_134690). Two of the 12 candidates (Smp_186930 and Smp_033950) were not annotated as TF in the KEGG search but indicated as orthologues to *C. elegans* counterparts. Based on a meta-analysis of gene expression for *S. mansoni* developmental stages^40^, the Stem/germinal cell cluster specific markers (**Figure 4A**) with promoter motifs and TFBSs identified above, tend to show a high expression in the ovary of mature females, miracidia and sporocysts (**Supplementary Figure S10A**). Interestingly, the *sma-4* ortholog Smp_033950 (Mothers against decapentaplegic or MAD homolog) also showed a high expression across these developmental stages, particularly in sporocysts, relative to the other stages (**Supplementary Figure S10A**). At the single-cell level, this gene was expressed across all cell clusters, with the highest expression and percentage of cells in the Neuron cluster (**Supplementary Figure S10B**). Even though the TF Smp_033950 was expressed in <5% of the stem/germinal cells (**Supplementary Figure S10B**), it was the only one among the 12 putative TFs identified in this study (**Supplementary Table S13**) that showed a Stem/germinal cluster-specific expression and almost exclusively in subclusters 1 and 2 (**Supplementary Figure S10C**). To reveal interactions between the TF Smp_033950 and its tentative target genes, we investigated the co-expression at the single cell level of this TF and the Stem/germinal cell markers predicted to have TFBSs for it, namely Smp_113620 and Smp_317550 with two predicted binding sites, and Smp_074390 with a single predicted site (**Figure 4C**). In addition, we included a representative stem/germinal cell-specific marker with no predicted TFBS for Smp_033950 (e.g., Smp_341240). No strong evidence of co-expression between the TF Smp_033950 and either of the stem/germinal marker genes was observed; for each of the analysed paired genes less than three cells showing a convincing co-expression signal were identified (**Supplementary Figure S11**). Moreover, analysis of genes not predicted to have TFBSs for the Smp_033950 revealed a similar number and pattern of co-expressing cells (e.g., Smp_341240; **Supplementary Figure S11**). However, the absence of evidence cannot at this stage be taken as evidence that there is no co-expression, given the high frequency of gene expression signal dropouts single-cell transcriptomic datasets^41^.

## Discussion

In this study, we employed single cell transcriptomics, followed by spatial validation, to unveil the cellular components of the mother sporocyst, the life cycle stage that marks the start of rapid asexual proliferation within the intermediate host. Due to the experimental challenges of studying parasites within snail tissues, we have used mother sporocysts cultured *in vitro* for 5 days, an approach that has been shown to mirror many aspects of *in-vivo* parasite development in snails^42–45^. For instance, the first scRNA-seq study in schistosomes described the transcriptomic signatures of proliferating cells derived from mother sporocysts transformed *in vitro*^19^. The authors validated their findings in parasites developing *in vivo*, following parasite progeny through genesis of cercariae and intra-mammalian stages^19^. Molecular and cellular differences cannot be completely ruled out; however, *in-vivo* validation of observations across several stages of development^19^ provides strong evidence that the *in-vitro* miracidium–sporocyst transformation, and culture of early sporocyst stages, is a reliable model to study the parasite *in-vivo* development.

Striking changes have been reported during the miracidium–sporocyst transition, including shedding of the ciliary plates, tegumental remodelling and an overall tissue reorganisation^39^ that transforms the free-living miracidium with ∼365 cells^18^ into a simple ‘sac-like structure’ enriched with stem/germinal cells^10^. With the assistance of a novel machine learning approach^24^, we estimated an average of 169 nuclei in a single D5 mother sporocyst (range of 112 to 254 nuclei), and have revealed previously uncharacterised variability in cell number among the parasites at this developmental stage. The variability in cell number was expected, as asynchrony has been described during both *in-vitro* and *in-vivo* development of this and other developmental stages of schistosomes^43,44,46^. The total number of sporocyst cells that passed the scRNA-seq QC-filtering steps was 601(i.e., >3.5-fold the number of cells estimated to be present in a single D5 mother sporocyst). Similar to our previous findings in early schistosomula, using the same dissociation and cell staining protocol^17^, we were, however, not able capture all cell types known to be present in this developmental stage, such as flame cells of the protonephridia system^47^. Limitations have been previously reported for the FDA/PI staining of live/dead cells, and for the cell sorting, including unspecific FDA signals in the cell suspension^48^, and fragile cells being destroyed or lost^49^, respectively. Optimised protocols to avoid dissociation-induced artefacts, and improve live/dead cell staining and cell sorting are being currently tested by us^18^ and others^50^.

Notwithstanding the above limitations, we were able to capture from whole mother sporocysts the transcriptomic signatures of most of the cell types previously identified by electron microscopy in the parasite up to ten days post-infection^6^. We identified six discrete cell clusters that showed distinct transcriptomic signatures, with consistent enriched GO term, and defined anatomic localisations in the whole animal, confirming previous electron microscopy findings^6,8,10,11^.

Interestingly, D5 sporocysts no longer show a well-defined brain structure, as described in the miracidium^18,51^. The complex nervous system described for miracidium at the single cell level, comprising >45% of cells^18^, appears to have degenerated into only a handful of cells in the D5 sporocyst. Approximately24 hours after infection, mother sporocysts newly-transformed *in vivo* already show neurons with evident degenerative changes^10^. The Muscle cluster marker *myosin heavy chain* was used to localise the myocytes that, as expected, colocalized with F-actin revealed by phalloidin stain. Muscle fibres also showed signs of degeneration, becoming thinner over time after the infection^10^. In spite of these degenerative changes in the neuromuscular system, sporocysts in culture are slightly motile and this activity increases when exposed to serotonin and serotonin receptor antagonists^52^. It has been reported that the infection of *Biomphalaria glabrata* snails with *S. mansoni* is associated with a reduction of serotonin and dopamine levels in the host brain^53^; however, the physiological role of these monoamine neurotransmitters on the sporocyst and its interaction with the host remains unknown.

The parenchyma cells identified in the D5 sporocyst showed a similar distribution to those in the miracidium^18^. Whilst these cells deliver energy, nutrients, and vitamins to the highly proliferating stem/germinal cells, they also start to degenerate and shrink to provide space for the early daughter sporocyst embryos^10,39^. Consistent with particles of complex carbohydrates and lipid droplets being described in the cytoplasm of parenchyma cells (historically described as ‘interstitial cells’)^10^.

Indeed, it has been suggested that the parenchyma cells in the mother sporocyst store metabolic energy for use by the highly-proliferative stem/germinal cells during the early development of daughter sporocyst embryos^6,10^. This hypothesis is supported by the observation of parenchyma cells degenerating and dying while daughter sporocysts developed within the brood chamber in the mother sporocyst^6^.

The tegument of the miracidium is remodelled within the first ∼5 hours of infecting a snail, a process that involves shedding the ciliary plates and expanding the syncytial ridges between them to cover the parasite surface^11^. Similar to the intra-mammalian developmental stages the sporocyst tegument is a syncytium connected to cell bodies below the muscle layer, which has prompted comparisons of miracidium-to-sporocyst and cercaria-to-schistosomulum transformation mechanisms^11^. Our FISH experiments confirmed that the tegument marker *MEG-6* is expressed on the surface of the parasite. The GO term analysis revealed unexpected evidence of high nucleotide/nucleotide metabolic activity in the tegument cells, including evidence of purine and pyrimidine salvage. These biological processes enriched in the sporocyst tegument may be specific to this developmental stage as none of these GO terms have been identified in the tegument of the miracidium^18^, schistosomulum^17^ or adult^16^. In addition, GO terms associated with microvilli (‘actin filament binding’ and ‘microtubule-based process’) were significantly enriched in the sporocyst tegument. In fact, the GO term ‘microtubule-based process’ was the only biological process in the tegument shared among the four analysed developmental stages. By day 3 post infection, the sporocyst is completely covered with numerous microvilli that increase the exchange surface of the tegument revealing high absorptive activities during the rapid development of the parasite^10,39^. Unlike the schistosome intra-mammalian developmental stages, the sporocysts lack a specialised digestive epithelium (i.e., gastrodermis). However, and similar to cestodes^54^ our findings suggest that the absorption and metabolism of snail-derived nutrients may be accomplished by the sporocyst tegument.

Schistosomiasis is a disease of stem cells, as these cells are the ultimate driver of life cycle propagation, development, and tissue homeostasis^55^. The stem cell system has been studied by both ‘bulk’-^20^ and single cell-transcriptomic^13,19^ approaches in schistosomes and their free-living relative, planaria^56^. Three discrete stem cell populations, named *kappa*, *delta* and *phi* (based on their key marker genes) have previously been identified in the mother sporocyst^13,19^. Furthermore, stem cell progenies were followed *in vivo* throughout the intra-snail and intra-mammalian developmental stages^4^. Using SAM, a recently developed algorithm that can find meaningful differences in systems with high intrinsic variability^30^, we identified three discrete Stem/germinal cell subclusters (named 0, 1 and 2), but were unable to unambiguously assign any of them to either *kappa*, *delta* or *phi* cell populations. There are, however, substantial technical differences between the two studies. In the present study, we used a dissociation protocol that enriched for all live cells and used 10X Genomics Chromium to capture single cells before sequencing. Wang *et al.* (2018) performed different dissociation — brief detergent treatment followed by trypsin digestion — then enriched EdU+ proliferative stem cells by FACS and captured them individually on a microfluidic RNAseq chip^19^ before sequencing. Although the stem-cell-targeted study only analysed 35 individual cells, it enabled deeper sequencing per cell, hence higher resolution to accurately discriminate *kappa*, *delta* and *phi* cells^19^. Sub-cluster 0, identified in our dataset, showed a significant enrichment for GO terms associated with ribosome activity, mRNA processing and protein translation. This is consistent with the observation of germ cell ribosomes being replaced by polysomes one day after infection, and suggests that the cells are preparing for division^10^. Based on expression of the discriminatory marker *nanos* in subclusters 0 and 1, both clusters have some semblance of being *kappa* and *delta* cells, whereas subcluster 2 may correspond to the *phi* cell population because it lacked *nanos* expression and was enriched with genes commonly associated with the tegument such as dynein light chain and tetraspanin. Five days after infection, daughter sporocyst embryos begin to be surrounded by a primitive epithelium^6^. It is therefore tempting to speculate that the Stem/germinal subcluster 2 cells may be associated with somatic/temporary larva structures such as daughter sporocyst primitive epithelium, as has previously been described for *phi* cells^13,19^. Further studies will address this hypothesis.

Single cell transcriptomics have revealed tissue-and cell-specific regulatory networks, transcription factors (TFs) and transcription factor binding sites (TFBSs) in model organisms^57^. Therefore, we decided to explore the enrichment of promoter motifs in marker genes specific for two key clusters: the Stem/germinal and Tegument-1 clusters. Most of the enriched promoter motifs in Stem/germinal-and Tegument-1-specific marker genes showed similarity to known transcription factor binding sites (TFBSs) in *C. elegans,* including sites for the homeobox genes *ceh-22*, *vab-7*, *unc-30* ^58^ and the forkhead/winged helix factor *pha-4*, ortholog of which (i.e. *S. mansoni foxA*) is expressed during the early development of the schistosomulum oesophageal gland^59^, but not in the sporocyst. With the assistance of bulk RNAseq metadata across several developmental stages^40^, we identified the schistosome orthologue of SMAD/NF-1 DNA-binding domain factor sma-4 (Smp_033950, *Smad4* in WBPS17), highly expressed in ovary of mature females, miracidia and sporocyst and for which TFBSs were present in the promoter region of Stem/germinal cell cluster marker genes. The Smad proteins belong to a family of intracellular transducers in the Transforming Growth Factor β (TGF-β) pathway involved in *S. mansoni* development, host-parasite interaction and male-female pairing^60,61^. *S. mansoni* Smad4 (SmSmad4) has previously been characterised^62^; following the binding of TGF-β/Activin ligand to TGF-β receptor, SmSmad1 and SmSmad2 (receptor-regulated Smads or R-Smads) interact with SmSmad 4 in the cytoplasm, which in turn phosphorylate the Smad-complex that is translocated to the nucleus to control the expression of target genes^61^. The relative expression of SmSmad1, 2 and 4 is stable across the intra-mammalian stages but, consistent with our own findings, SmSmad4 is overexpressed relative to the other two molecular partners in the egg and mother sporocyst^62^. This would suggest a specific role during the development of intra-snail stages or interaction with the snail. The TGF-β pathway plays key roles in cell differentiation, proliferation and apoptosis, all critical processes for sporocyst development. Here, we have identified the cell clusters where SmSmad 4 is expressed and its tentative promoter binding sites in target genes.

This study represents the first molecular characterisation of both somatic and germinal cells of the schistosome mother sporocyst. This developmental stage is essential for the parasite amplification that in turn facilitates a successful propagation of the life cycle and sustains the high prevalence of the infectious disease in endemic areas. A better understanding of the mechanisms underlying these processes could lead to novel control strategies for schistosomiasis and related trematodiases.

## Data (and Software) Availability

All raw sequence data is deposited in the ENA under the project accession ERP137194 and sample accession numbers: ERS11891013, ERS11891016, ERS11891015, ERS11891014 (**Supplementary Table S1**). For the purpose of Open Access, the authors have applied a CC BY public copyright licence to any Author Accepted Manuscript version arising from this submission.

## Author Contributions

Conceptualization: CD, GR; Data Curation: CD, TA, ZL; Formal Analysis: CD, TA, ZL, TGRA, KAR, GR; Funding Acquisition: MB; Investigation: CD, JG, CH, ST, TGRA, GR; Methodology Development: CD, TA, ZL, TGRA, KR, GR; Project Administration: MB, GR; Resources: CD, TA, GR; Software Programming: CD, TA, ZL; Supervision: MB, GR; Validation: TA, KAR, GR; Visualization: CD, TA, ZL, GR; Writing – Original Draft Preparation: GR; Writing – Review & Editing Preparation: all authors

## Acknowledgements

We thank our colleagues at the Wellcome Sanger Institute (WSI): Simon Clare, Cordelia Brandt, Catherine McCarthy, Katherine Harcourt and Lisa Seymour for assistance and technical support with animal infections and maintenance of the *Schistosoma mansoni* life cycle; Geetha Sankaranarayanan and Magda Lotkowska for expert technical support; Sarah Buddenborg and Stephen R. Doyle for providing access to the latest versions of the *S. mansoni* genome assembly; David Goulding and Claire Cormie at the Electron and Advanced Light Microscopy facility for technical support with confocal microscopy; Nancy Holroyd, Mandy Sanders and Siobhan Austin-Guest for project and sequencing coordination. We also acknowledge Andrew Gillis from the Zoology Department, University of Cambridge for producing the image of a representative D5 mother sporocyst shown in Figure 1A. This research was funded in part by the Wellcome Trust [Grant numbers 206194 and 107475/Z/15/Z] and a UKRI Future Leaders Fellowship awarded to GR [Grant number MR/W013568/1].

## Supplementary Information

### Supplementary Figures and Tables

**Supplementary Figure S1.**
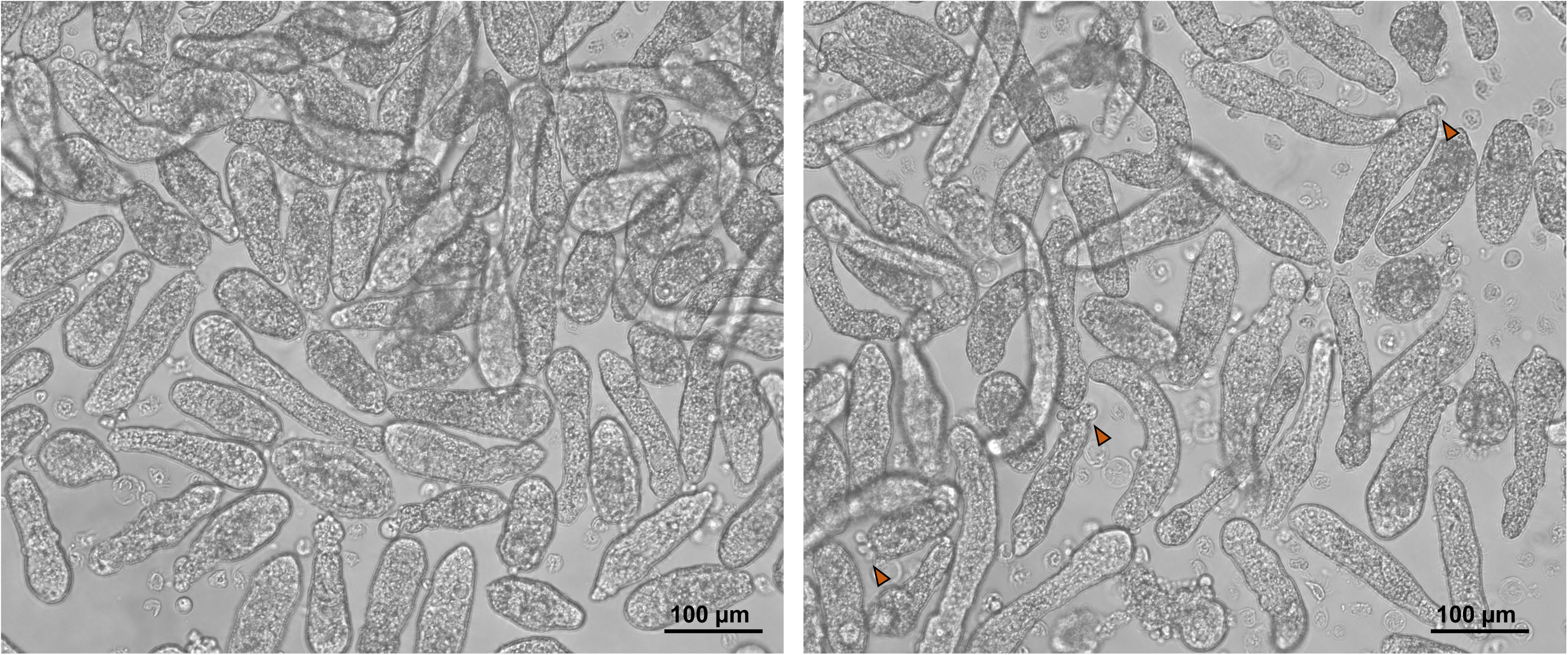
Live D5 mother sporocysts before processing for single cell RNA-seq or FISH. Representative pictures of live *S. mansoni* mother sporocysts transformed from freshly-hatched miracidia and cultured for 5 days in sporocyst medium under hypoxia at 28 °C for 5 days. More than 95% of the parasites have fully transformed into mother sporocysts, and only a minority of parasites showed 1-3 cilia plates still attached to either the anterior or posterior pole (orange arrowheads). Scale bar: 100μm.

**Supplementary Figure S2.**
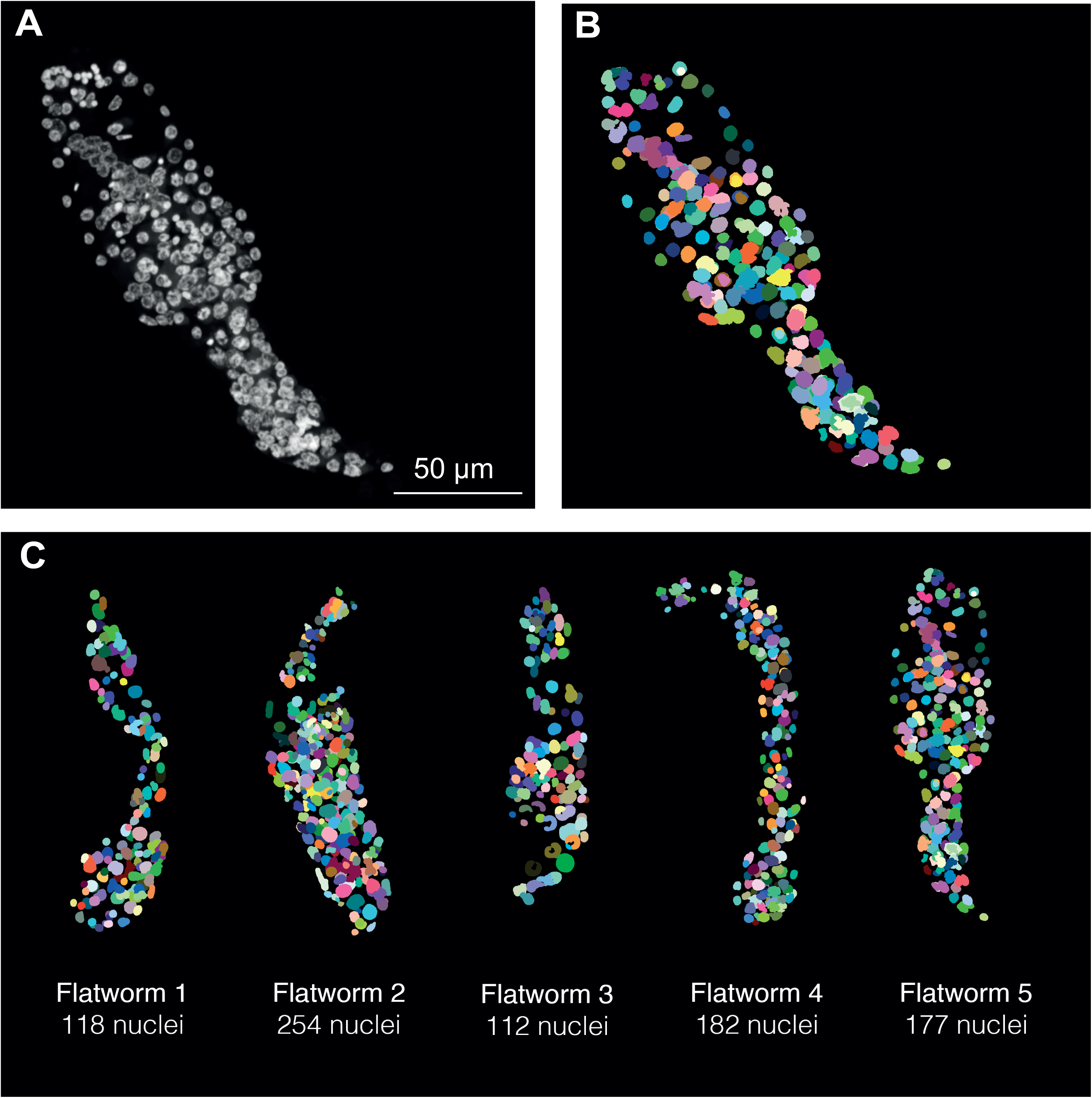
Nuclear segmentation of D5 mother sporocysts. Nuclear segmentation of a representative D5 sporocyst used to generate the training data samples: pre-processed DAPI signal (**A**) and final segmentation map (**B**). **C.** Nuclear segmentation of 5 parasites revealed a total number of nuclei ranging from 112 to 254, with a mean of 168.6 and standard deviation of 57.68. Scale bar: 50μm.

**Supplementary Figure S3.**
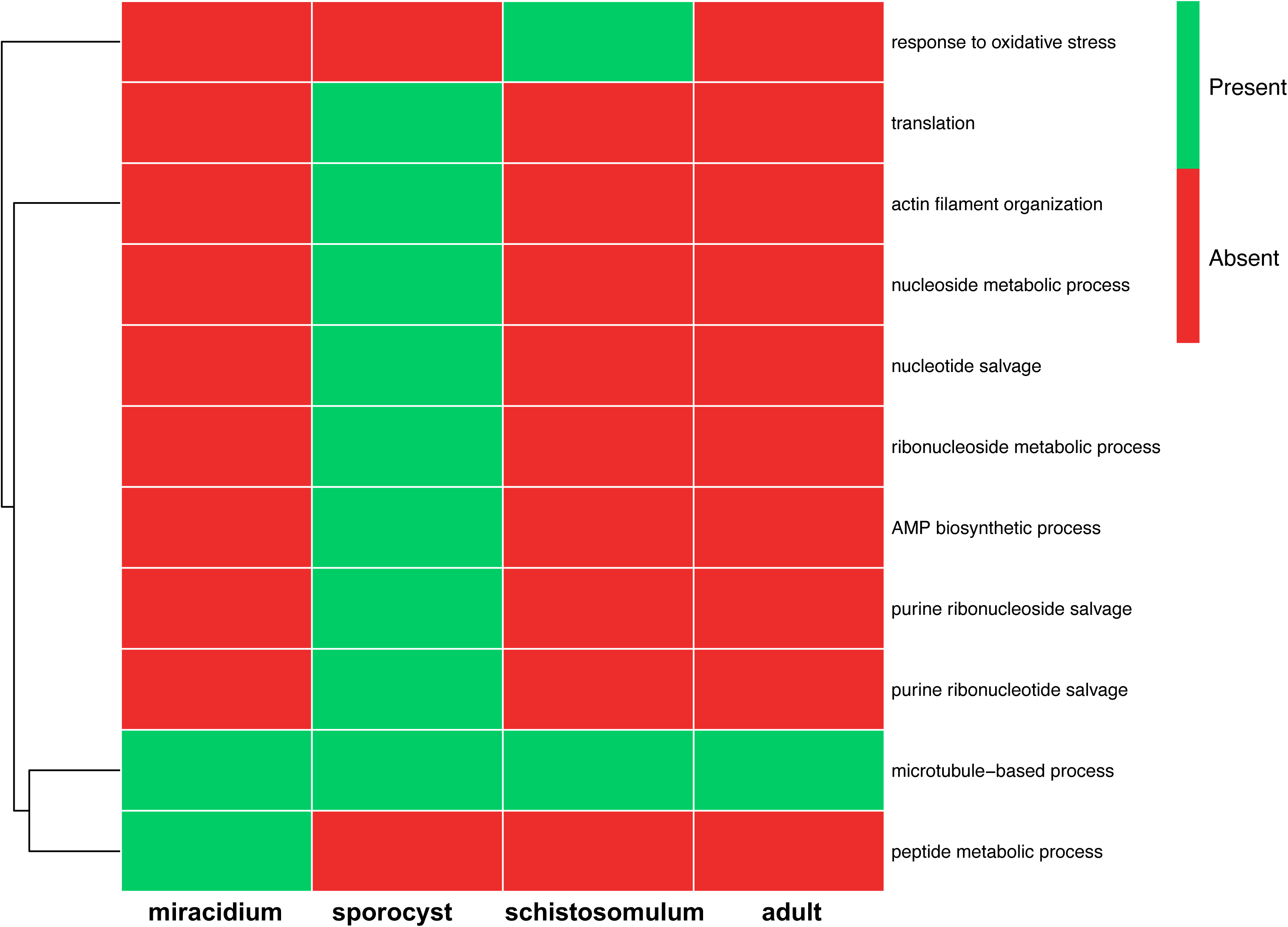
Biological processes in the tegument of *S. mansoni* developmental stages. Clustered heatmap showing presence (green)/absence (red) of the tegument GO terms (Biological Processes) in the miracidia, sporocyst, schistosomula and adult worms, as indicated.

**Supplementary Figure S4.**
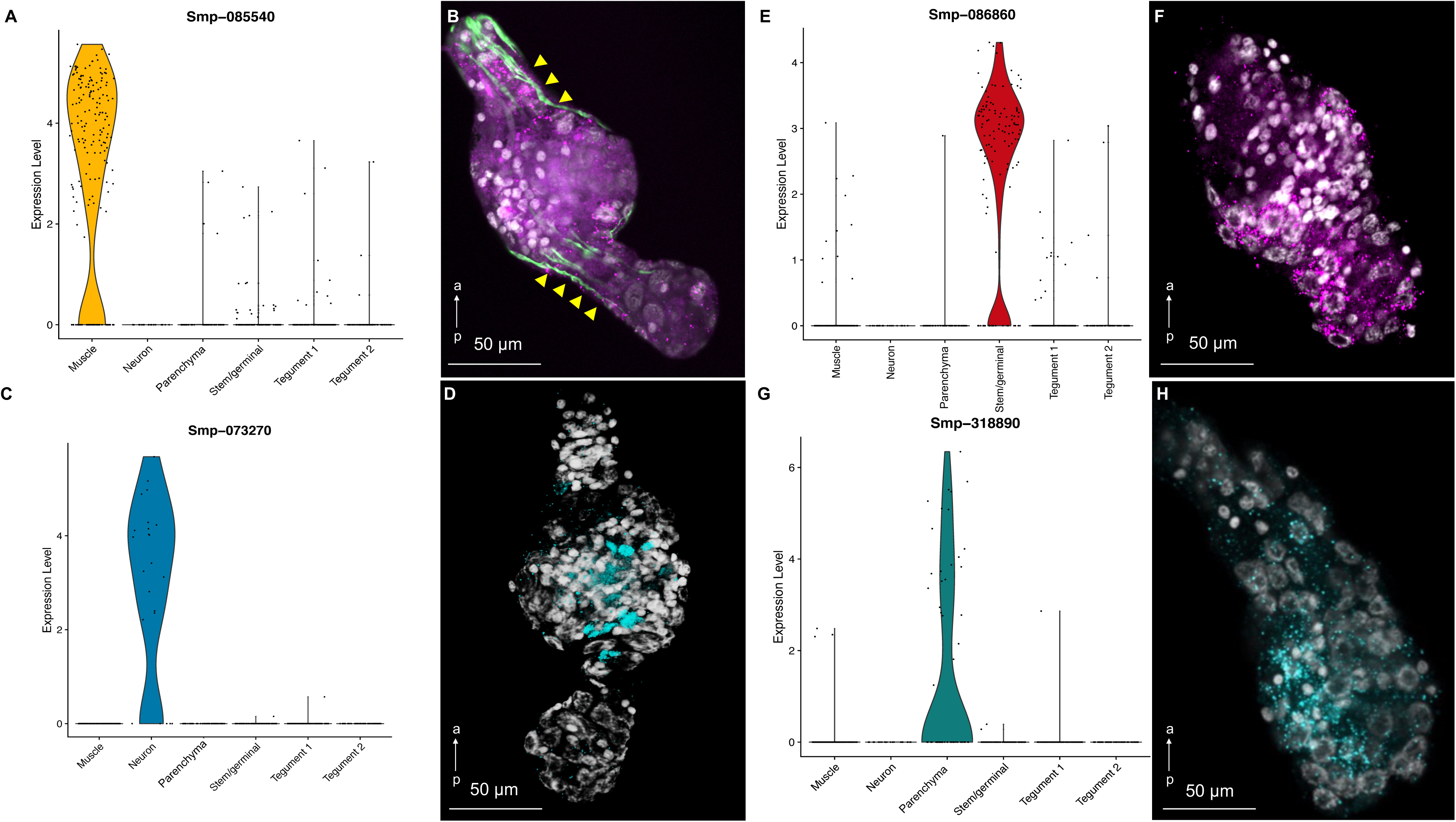
Spatial validation of cell clusters. **A.** Violin plot showing the expression level of the muscle marker *myosin heavy chain* (Smp_085540) across the six cell clusters. **B.** Muscle cells expressing *myosin heavy chain* (*S*mp_085540; magenta) revealed by FISH, co-localised with phalloidin-stained actin filaments (green). Yellow arrowheads indicate co-localisation of *myosin heavy chain* and actin filaments. **C.** Violin plot showing the expression level of the neuron marker *neuroendocrine protein 7b2* (Smp_073270) across the six cell clusters. **D.** Neuron cluster cells expressing neuroendocrine *protein 7b2* (Smp_073270; cyan) revealed by FISH (same specimen as the one shown in Figure 2B). **E.** Violin plot showing the expression level of the stem/germinal marker *histone H2A* (Smp_086860) across the six cell clusters. **F.** Stem/germinal cells expressing *histone H2A* (Smp_086860; magenta) revealed by FISH. **G.** Violin plot showing the expression level of the parenchymal marker *hypothetical protein* (Smp_318890) across the six cell clusters. **H.** Parenchyma cells expressing *hypothetical protein* (*Smp_318890*; cyan) revealed by FISH. DAPI staining in grey. Scale bar: 50μm, a←p: anterior-posterior axis (Panels B, D, F, H)

**Supplementary Figure S5.**
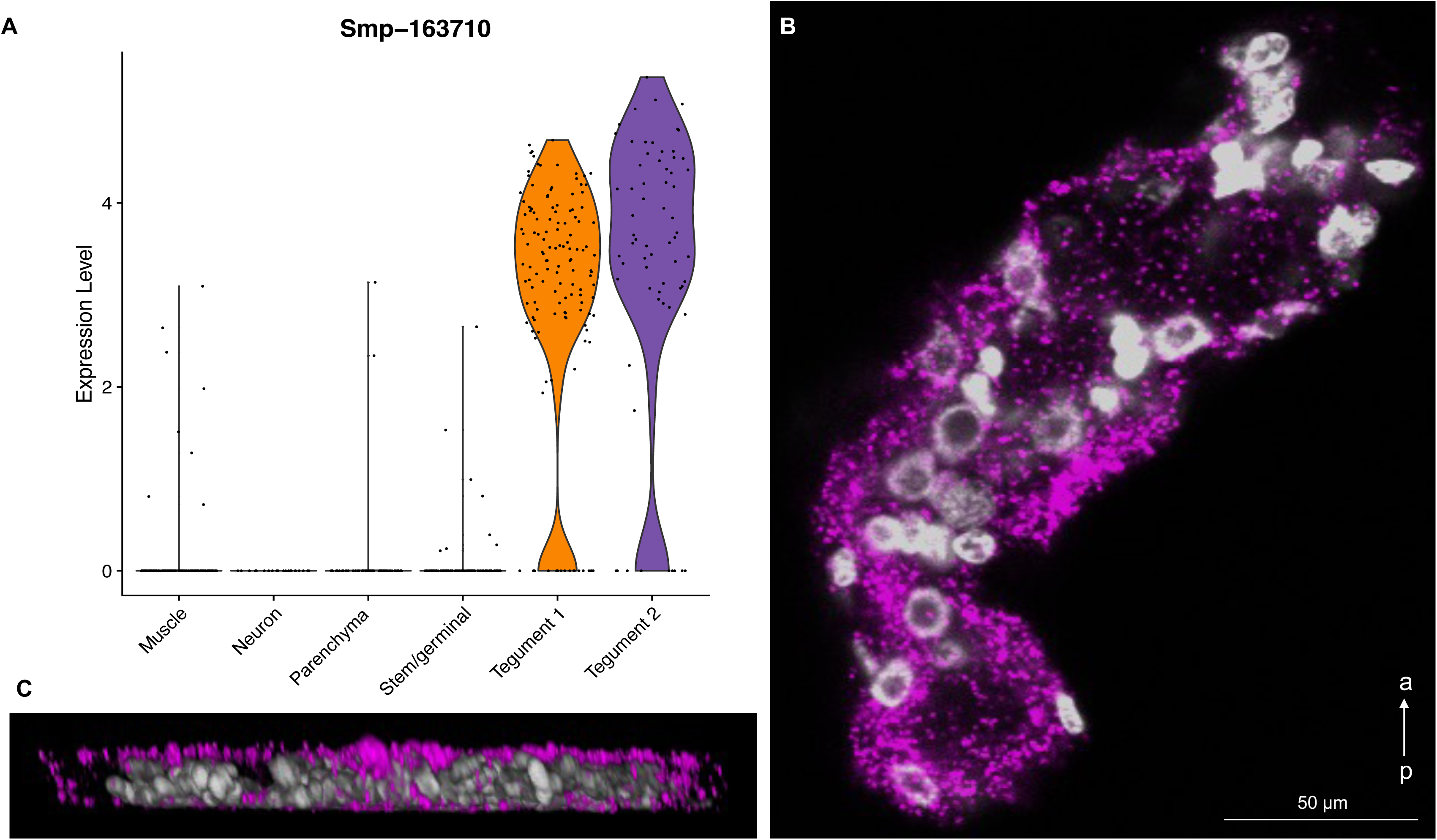
Spatial validation of the tegument clusters. **A.** Violin plot showing the expression level of the pan-tegumental micro-exon gene 6 or *MEG-6* (Smp_163710) across the 6 cell clusters. **B.** FISH of micro-exon gene 6 or *MEG-6* (Smp_163710)-expressing cells (magenta) identified the tegument clusters. DAPI staining in grey. Scale bar: 50μm, a←p: anterior-posterior axis. **C.** Sagittal projection of a parasite showing the tegumental expression of *MEG-6*. DAPI staining in grey.

**Supplementary Figure S6.**
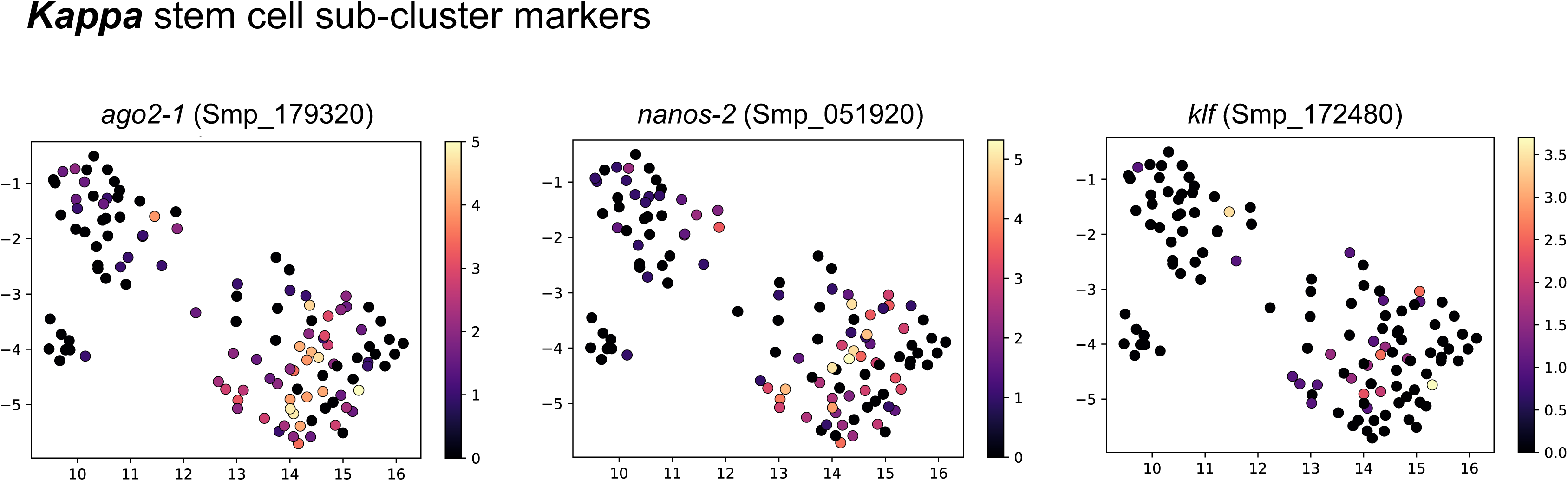
*Kappa* cell marker genes. Uniform Manifold Approximation and Projection (UMAP) representation of 119 stem/germinal cells clustered using self-assembling manifold (SAM) algorithm. Single-cell expression level of the 3 indicated marker genes for *kappa* cells. The average gene expression level for each marker is represented by a colour gradient from dark blue (low expression) to bright yellow (high expression).

**Supplementary Figure S7.**
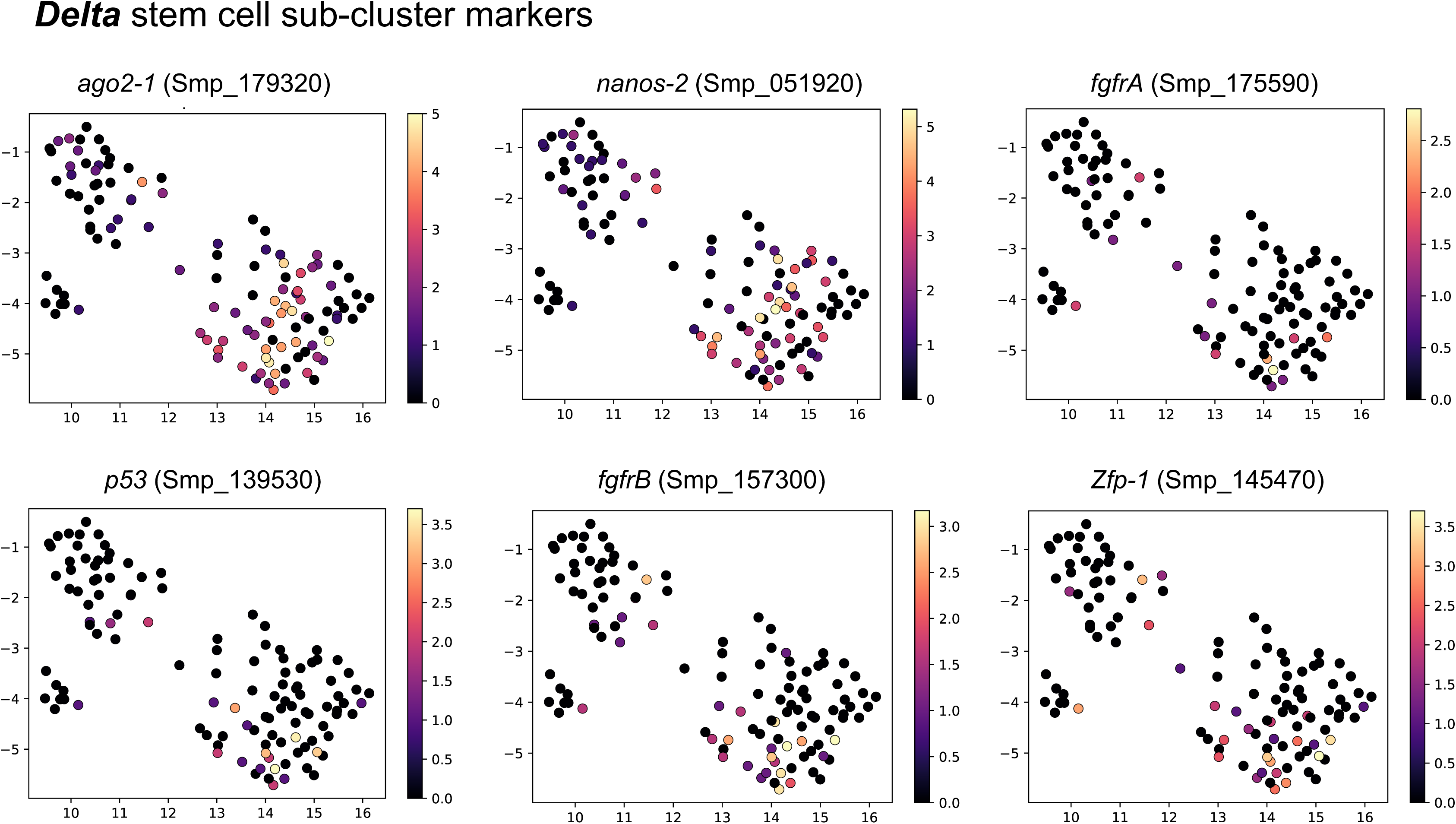
*Delta* cell marker genes. Uniform Manifold Approximation and Projection (UMAP) representation of 119 stem/germinal cells clustered using self-assembling manifold (SAM) algorithm. Single-cell expression level of the 6 indicated marker genes for *delta* cells. The average gene expression level for each marker is represented by a colour gradient from dark blue (low expression) to bright yellow (high expression).

**Supplementary Figure S8.**
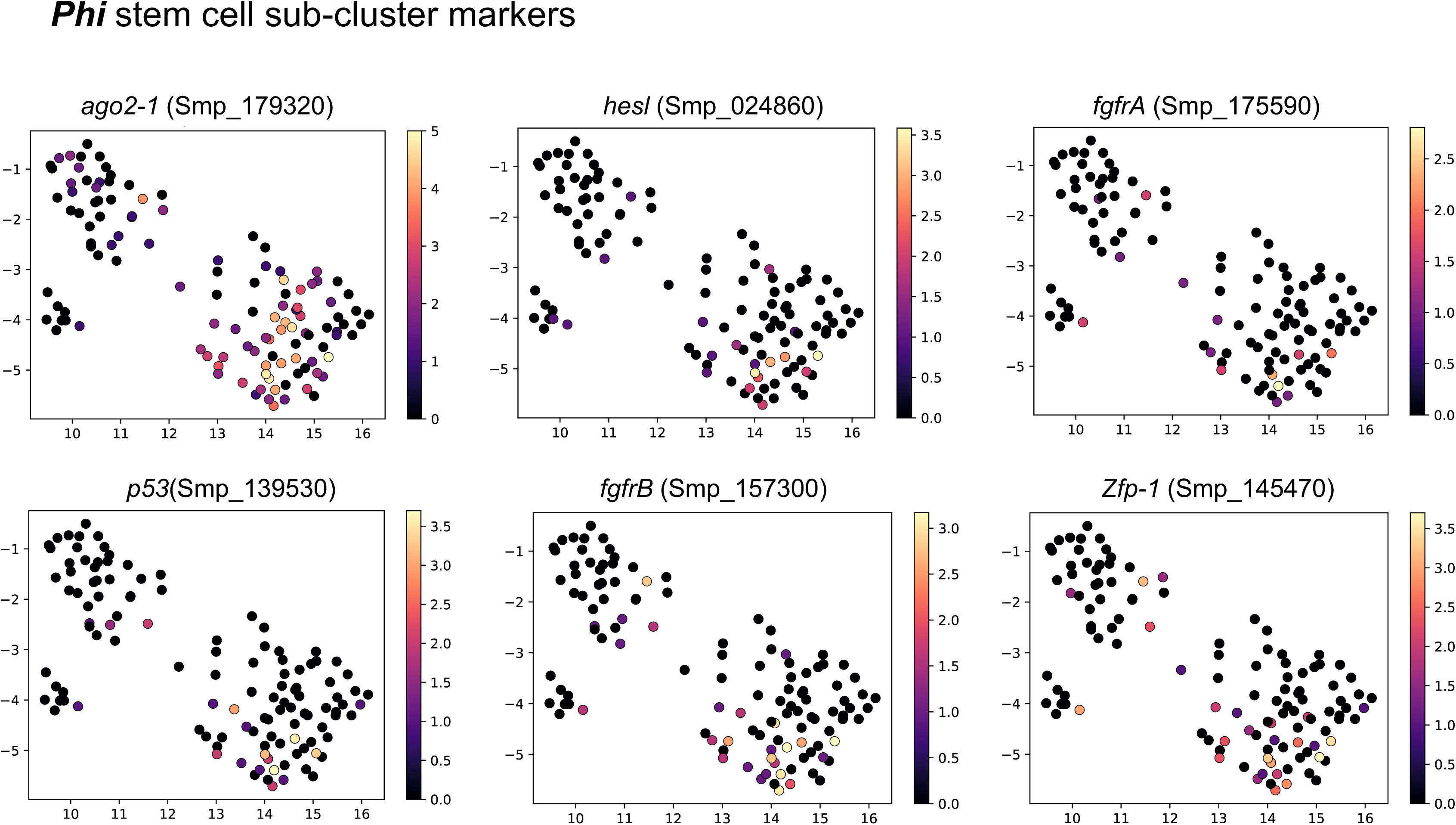
*Phi* cell marker genes. Uniform Manifold Approximation and Projection (UMAP) representation of 119 stem/germinal cells clustered using self-assembling manifold (SAM) algorithm. Single-cell expression level of the 6 indicated marker genes for *phi* cells. The average gene expression level for each marker is represented by a colour gradient from dark blue (low expression) to bright yellow (high expression).

**Supplementary Figure S9.**
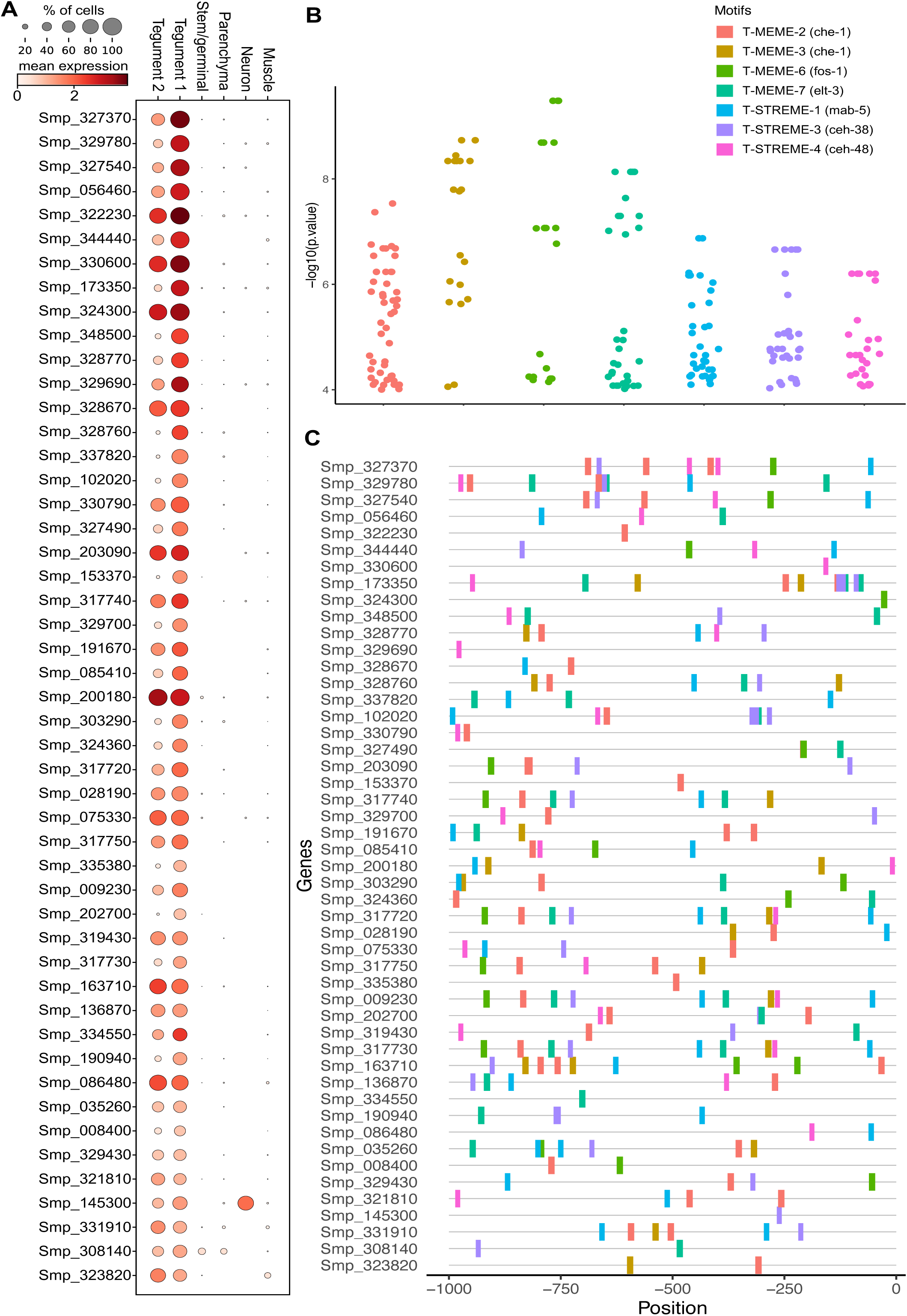
Promoter motif and transcription factor binding sites in tegument cells. **A.** Dot plot showing the expression level of the 49 Tegument-1 cell cluster-specific marker genes used for the analysis. Fraction of cells (%) and mean expression are indicated. The average gene expression level for each marker is represented by a colour gradient from white (low expression) to dark red (high expression). **B**. Distribution of the –log10(p values) for the top 10 ranked motifs identified in the 49 Tegument-1 cell marker genes. The x-axis indicates motif names from XSTREME and significant match (p < 0.05) to known Transcription Factors Binding Sites (TFBSs) in the JASPAR 2022 nematode dataset (https://jaspar.genereg.net/downloads/). Seven of the 10 are shown, as they are the ones which showed matched jaspar nematode TFBS (see **Supplementary Table S13**, motif shown in red). The y-axis represents log-transformed p values of each motif site shown in C. **C.** Predicted position distribution of the top 7 ranked motifs with matched TFBSs along the promoter region of the Tegument-1 cell marker genes. The promoter region was taken as 1 kb upstream of the Transcription Start Site (TSS). Full data provided in **Supplementary Table S14**.

**Supplementary Figure S10.**
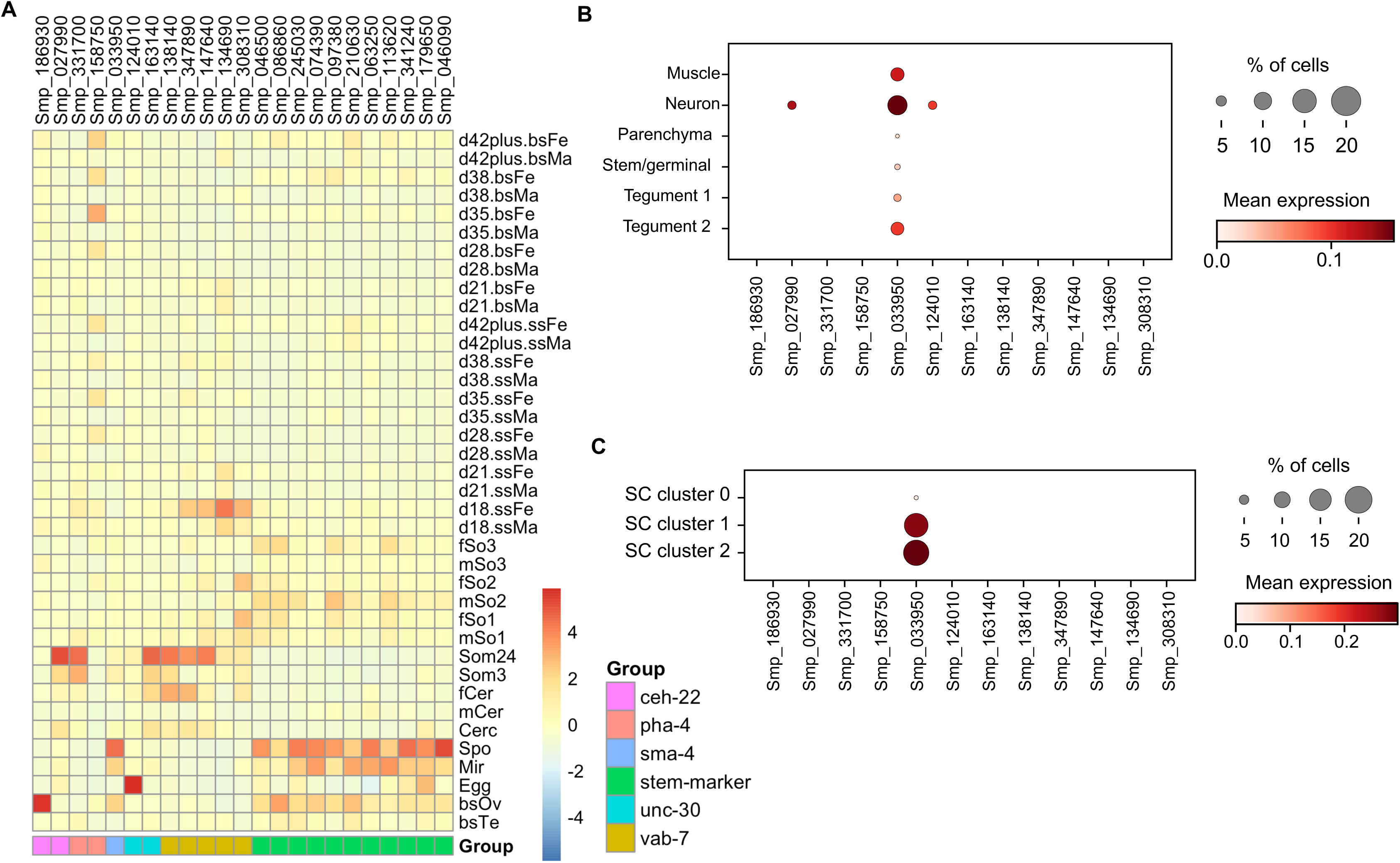
Bulk and single-cell expression of predicted transcription factors. **A.** Heatmap of relative expression of the Stem/germinal cell cluster marker genes that show within the promoter region tentative Transcription Factor Binding Sites (TFBSs) (indicated as stem-marker), and predicted TFs (i.e, *S. mansoni* genes orthologous to the *C. elegans* genes *ceh-22*, *pha-4*, *sma-4*, *unc-30*, and *vab-7* as indicated) across all developmental stages. ‘Bulk’ RNAseq data across all the indicated developmental stages were obtained from published data as described above. The average gene expression level for each marker is represented by a colour gradient from dark blue (low expression) to dark red (high expression). **B.** Dot plot depicting the fraction of cells (%) and mean expression of the predicted TFs at the single cell level across all the cell clusters. The average gene expression level for each marker is represented by a colour gradient from white (low expression) to dark red (high expression). **C.** Dot plot depicting the fraction of cells (%) and mean expression of the predicted TFs at the single cell level across the three stem/germinal sub-clusters. The average gene expression level for each marker is represented by a colour gradient from white (low expression) to dark red (high expression). SC: sub-cluster

**Supplementary Figure S11.**
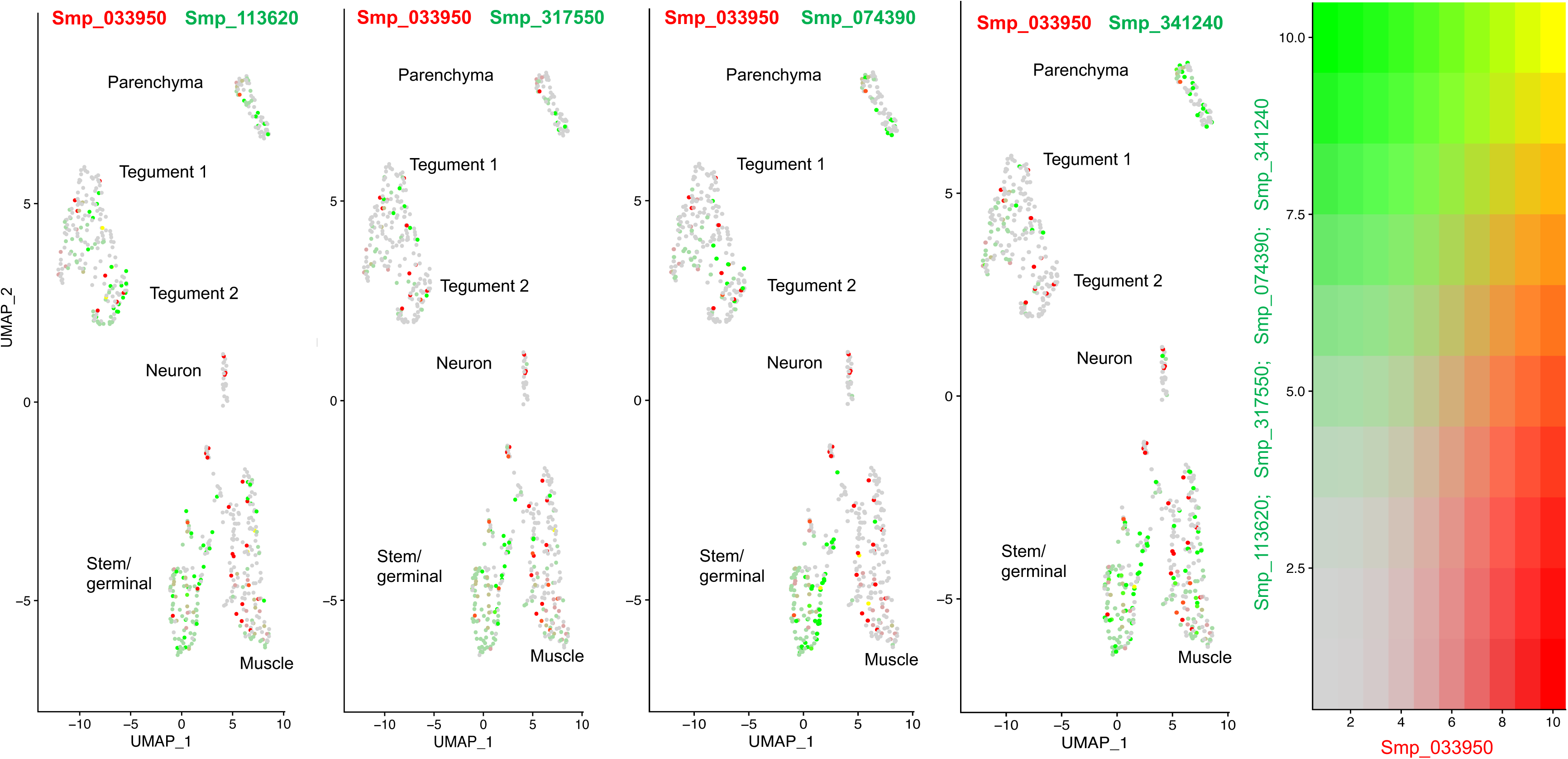
Co-expression analysis at the single cell level. Uniform Manifold Approximation and Projection (UMAP) representation of all cell clusters as indicated. Co-expression level of the TF Smp_033950 (in red) with each of the four Stem/germinal cell cluster markers (in green) predicted to have that two (Smp_113950, Smp_317550), one (Smp_074390) or no (Smp_341240) predicted binding site(s) for the TF Smp_033950. Bi-coloured expression panel for the TF Smp_033950 (red) and tentative target genes (green) is shown on the right. Yellow cells indicate high co-expression levels for both genes.

**Supplementary Table S1**. Samples and single cell RNA-Seq mapping statistics.

**Supplementary Table S2**. *In situ* hybridization probes.

**Supplementary Table S3**. Seurat marker genes in all cell clusters.

**Supplementary Table S4**. GO enrichment terms AUC=0.7, no specificity filtering applied (summarised in Figure 1C).

**Supplementary Table S5**. GO enrichment terms AUC=0.6, no specificity filtering applied.

**Supplementary Table S6**. Top 5 marker genes for each cell cluster (shown in Figure 2A).

**Supplementary Table S7**. SAM topology marker genes for all cells.

**Supplementary Table S8**. SAM topology marker genes for cells in the Stem/germinal cell cluster.

**Supplementary Table S9**. Top marker genes identified by SCANPY in the three SAM Stem/germinal cell subclusters.

**Supplementary Table S10**. All enriched STRING terms for the top 50 markers for each sub-cluster; minimum required interaction score of 0.9.

**Supplementary Table S11**. GO enrichment terms for stem/germinal subclusters AUC=0.7, no specificity filtering applied (summarised in Figure 3D).

**Supplementary Table S12**. xstreme_output_summary.

**Supplementary Table S13**. Summary-motif-tfbs.

**Supplementary Table S14**. ClusterMarker_Motif-TFBS.

**Supplementary Table S15**. NematodeTF_tophits

